# Visualizing pyrazinamide action by live single cell imaging of phagosome acidification and *Mycobacterium tuberculosis* pH homeostasis

**DOI:** 10.1101/2021.12.20.473466

**Authors:** Pierre Santucci, Beren Aylan, Laure Botella, Elliott M. Bernard, Claudio Bussi, Enrica Pellegrino, Natalia Athanasiadi, Maximiliano G. Gutierrez

**Affiliations:** Host-Pathogen Interactions in Tuberculosis Laboratory, The Francis Crick Institute, 1 Midland Road, London, NW1 1AT, United Kingdom

**Keywords:** Tuberculosis, Microenvironments, Antibiotics, Intracellular Pharmacokinetics, Human Macrophages

## Abstract

The intracellular population of *Mycobacterium tuberculosis* (Mtb) is dynamically segregated within multiple subcellular niches with different biochemical and biophysical properties that, upon treatment, may impact antibiotic distribution, accumulation, and efficacy. However, it remains unclear whether fluctuating intracellular microenvironments alter mycobacterial homeostasis and contribute to antibiotic enrichment and efficacy. Here, we describe a dual-imaging approach that allows quantitative monitoring of host subcellular acidification and Mtb intrabacterial pH profiles by live-fluorescence microscopy in a biosafety level 3 laboratory. By combining this live imaging approach with pharmacological and genetic perturbations, we show that Mtb can maintain its intracellular pH independently of the surrounding pH in primary human macrophages. Importantly, we show that unlike bedaquiline (BDQ), isoniazid (INH) or rifampicin (RIF), the front-line drug pyrazinamide (PZA) displays antibacterial efficacy by acting as protonophore which disrupts intrabacterial pH homeostasis *in cellulo*. By using Mtb mutants with different intra-macrophage localisation, we confirmed that intracellular acidification is a prerequisite for PZA efficacy *in cellulo*. We anticipate this dual imaging approach will be useful to identify host cellular environments that affect antibiotic efficacy against intracellular pathogens.

**Highlights:**

- Mtb maintains its intrabacterial pH inside both acidic and neutral subcellular microenvironments of human macrophages
- Pyrazinamide, but not other frontline antibiotics, acts as a protonophore *in cellulo*
- Pyrazinamide-mediated intrabacterial pH homeostasis disruption and antibacterial efficacy requires host endolysosomal acidification
- Cytosolic localisation mediated by ESX-1 contributes to pyrazinamide antibacterial activity resistance
- Pyrazinamide conversion into pyrazinoic acid by the pyrazinamidase/nicotinamidase PncA is essential for its protonophore activity and efficacy *in cellulo*

## Introduction

Tuberculosis (TB), caused by *Mycobacterium tuberculosis* (Mtb), remains one of the deadliest infectious disease worldwide (WHO, 2021). In 2020, it was estimated that almost 10 million people developed the active form of the disease and 1.3 million people died from TB, and recent data indicate that the COVID-19 pandemic is disrupting access to TB care and treatment (WHO, 2021).

Drug-susceptible TB treatment relies on a standard chemotherapy regimen that includes four first-line antibiotics, rifampicin (RIF), isoniazid (INH), ethambutol (EMB) and pyrazinamide (PZA), which are administered over a period of at least six months (WHO, 2021). This extensive treatment is often associated with side-effects and toxicity affecting compliance and contributing to the emergence of antibiotic resistant strains (Gulbay et al., 2006). In that context, it is crucial to better understand how current anti-TB chemotherapies work to develop a new generation of efficient, fast-acting, and compliance-friendly treatments.

TB is a complex disease in which Mtb infection is mainly characterized by the formation of heterogeneous pulmonary granulomatous lesions that evolve dynamically over time (Cadena et al., 2017; Lenaerts et al., 2015). Inside these highly structured cellular aggregates, macrophages constitute the primary niche used by the tubercle bacilli to survive, replicate and disseminate (Cohen et al., 2018). To survive and replicate within host macrophages, Mtb has successfully developed multiple strategies to counteract host cell defence mechanisms (Bussi and Gutierrez, 2019). Among them, both modulation and subversion of phagosome maturation and its ability to survive within acidic and hydrolytic microenvironments have been demonstrated to be crucial for the intracellular lifestyle of Mtb (Brodin et al., 2010; Levitte et al., 2016; MacGurn and Cox, 2007; Pethe et al., 2004; Stewart et al., 2005; Vandal et al., 2008). In addition, Mtb can damage the phagosome through the action of its type VII secretion system ESX-1 and cell wall-associated phthiocerol dimycocerosates (PDIM) lipids to access the pH-neutral, nutrient-rich cytosol (Augenstreich et al., 2017; Barczak et al., 2017; Bernard et al., 2020; Lerner et al., 2017; Lerner et al., 2018; Quigley et al., 2017; van der Wel et al., 2007).

Because Mtb is localised in several intracellular niches, the generation of new drug regimens should consider the efficient targeting of Mtb intracellular populations residing within host cells. Ideally, anti-TB chemotherapy must include antibiotics with pharmacokinetic properties that allow agents to (i) penetrate lung tissue and infected cells, (ii) reach the wide-ranging subcellular compartments in which Mtb resides and (iii) be active in these specific microenvironments to finally display optimal antibacterial efficacy. Understanding how subcellular environments affect bacterial fitness (Huang et al., 2019; Rohde et al., 2007) but more importantly antibiotic localisation, exposure, and consequently efficacy against intracellular pathogens is crucial and have only recently begun to be investigated (Liu et al., 2016; Santucci et al., 2021).

Cell compartment specific consideration of bioactivity is of particular importance for antibiotics such as the front-line drug PZA, which was demonstrated to be highly potent against the tubercle bacilli within infected mice but didn’t display activity in standard culture conditions *in vitro* (Malone et al., 1952; Mc and Tompsett, 1954; Solotorovsky et al., 1952; Tarshis and Weed, 1953). Indeed, *in vitro* PZA requires an acidic pH below 5.5 to be effective against Mtb (Mc and Tompsett, 1954; Zhang et al., 2002; Zhang et al., 1999). In this widely accepted pH-dependent mechanism of action, PZA enters the bacteria by diffusion and is converted by the PncA enzyme to form the deprotonated negatively charged POA^(-)^ anion (POA^(-)^). This POA^-^ anion is then actively exported into the extracellular milieu. If Mtb faces an acidic environment, POA^-^ acquires a hydrogen cation to form the neutral protonated HPOA molecule (HPOA). This protonated form is able to diffuse across the bacterial envelope to finally disrupt intrabacterial pH homeostasis and membrane potential (Zhang et al., 2013). Several independent studies proposed that PZA/POA molecules act as pH-dependent protonophores *in vitro* to disrupt Mtb intrabacterial pH homeostasis (Darby et al., 2013; Fontes et al., 2020; Zhang et al., 1999; Zhang et al., 2003). However, this proposed pH-dependent mode of action of PZA/POA, where molecules acidify Mtb cytoplasm has been challenged (den Hertog et al., 2016; Peterson et al., 2015). Alternative pH-independent mechanisms underlying PZA/POA efficacy have been proposed (Gopal et al., 2019; Lamont et al., 2020) whereby POA^(-)^ acting at neutral pH can competitively inhibit the aspartate decarboxylase PanD and enhance its targeted degradation *via* the Clp system thus impairing the biosynthesis of Co-enzyme A (Gopal et al., 2020; Gopal et al., 2016; Sun et al., 2020). In this model, PZA/POA molecules are active regardless of the environmental pH surrounding Mtb and do not act as protonophores. Overall, it is notable that despite being used as a front line drug for almost 70 years in the clinic, the molecular and cellular mechanisms underlying PZA efficacy remain unclear (Lamont et al., 2020).

By using correlative light electron ion microscopy (CLEIM) approaches we showed that PZA/POA molecules require phagosomal residency and acidification to efficiently accumulate within Mtb and display optimal activity within human macrophages (Santucci et al., 2021). Mtb can transit through neutral and acidic environments multiple times during the infection cycle (Bernard et al., 2020; Schnettger et al., 2017), however it remains unclear how these spatial and temporal changes affect the intracellular activity of PZA and its impacts on intrabacterial pH homeostasis.

To address these questions, we developed a dual-imaging approach that allows monitoring of endolysosomal acidification and Mtb intrabacterial pH homeostasis in real time. Single cell quantitative analysis shows that Mtb can maintain its intrabacterial pH independently of the host pH. By live-cell imaging and tracking of Mtb bacterial populations residing within acidic endolysosomes, we show that long-term residence within acidic microenvironments, i.e. for several hours, is not sufficient to impact Mtb pH homeostasis. Using this approach, we describe the spatiotemporal dynamics of PZA mode of action within Mtb-infected human macrophages, showing that phagosomal restriction and host subcellular acidification are crucial for PZA/POA antibacterial efficacy. Finally, by using mycobacterial mutants with different phenotypes and intracellular lifestyles, we define how both host-driven and bacterial factors contribute to PZA efficacy in human macrophages.

## Results

### A live dual-imaging approach to monitor organelle and Mtb acidification

To concomitantly monitor macrophage organelle and Mtb acidification status, we developed a high-content dual-imaging approach that allows live-fluorescence microscopy visualization of infected human macrophages, including quantitative measurements of subcellular acidification and Mtb intrabacterial pH homeostasis. We first generated a Mtb reporter strain (Mtb pH-GFP) that constitutively produces a ratiometric pH-GFP indicator to dynamically record intrabacterial pH fluctuations *in vitro* and *in cellulo*, as previously described (Darby et al., 2013; Vandal et al., 2008). This reporter possesses two excitation maxima at wavelengths 405 nm and 488 nm and the ratio of 510 nm fluorescence emissions generated by excitation at these two excitation wavelengths varies as a function of the protonation state of the GFP fluorophore. Therefore, a lower 405 nm/488 nm ratio indicates a lower intrabacterial pH (Darby et al., 2013; Vandal et al., 2008). Then, human monocyte-derived macrophages (MDM) and human-induced pluripotent stem cell-derived macrophages (iPSDM) were infected with the reporter strain for 24 h, a time that allows bacteria to adapt intracellularly and localise in multiple niches (Bernard et al., 2020; Lerner et al., 2017). Infected cells were left untreated or pulsed for an additional 24 h with Concanamycin A (ConA), a selective v-ATPase inhibitor that inhibits endolysosomal acidification (Huss et al., 2002), and further stained with the lysosomotropic fluorescent probe LysoTracker to visualise acidic endolysosomes and determine host subcellular acidification profile. Next, Mtb-associated LysoTracker intensity and Mtb intrabacterial pH were analysed by automated high-content microscopy (**Figure S1**). A quantitative analysis in Mtb-infected MDM (median_CTRL_ = 422.6; IQR_CTRL_ = 334.9 and median_ConA_ = 241.6; IQR_ConA_ = 64.5, respectively) and Mtb-infected iPSDM (median_CTRL_ = 1964.2, IQR_CTRL_ = 1006.0 and median_ConA_ = 929.3; IQR_ConA_ = 1009.4, respectively) showed that the median Mtb-associated LysoTracker intensity was reduced by approximately 2-fold upon ConA treatment (**Figure 1A and Figure 1C**), confirming that endolysosomal acidification was impaired (Santucci et al., 2021). On the other hand, a quantitative analysis of Mtb intrabacterial pH in control or ConA-treated conditions were similar in both infected MDM (**Figure 1B**) and infected iPSDM (**Figure 1D**) with absolute median differences that were almost null (Δmedian_pH-GFP_ = 0.035 and 0.016 respectively), suggesting that intracellular acidification does not impact Mtb intrabacterial pH in human macrophages. To confirm that Mtb can maintain its intrabacterial pH independently of macrophage pH, we determined the Spearman’s correlation coefficient between Mtb pH-GFP ratio values with their corresponding associated LysoTracker intensity. There was no positive or negative association with correlation coefficients of *r*_*s*_ = 0.022, *p* < 0.01 and *r*_*s*_ = 0.062, *p* < 0.0001 in MDM or iPSDM respectively (**Figure 1E-1H**). Next, we performed live-image acquisition of Mtb-infected LysoTracker-stained iPSDM at higher-resolution (Bernard et al., 2020; Schnettger et al., 2017). In agreement with the previous findings (**Figure 1G and Figure 1H**), analysis of Spearman’s correlation coefficient did not show any correlation between subcellular acidification profile and Mtb intrabacterial pH with a value of *r*_*s*_ = -0.19, *p* < 0.0001 (**Figure S2**). Altogether these data support the notion that Mtb can maintain its own pH when facing a wide range of *in vitro* and *in cellulo* environmental pH (Darby et al., 2013; Fontes et al., 2020; Vandal et al., 2008).

**Figure 1:**
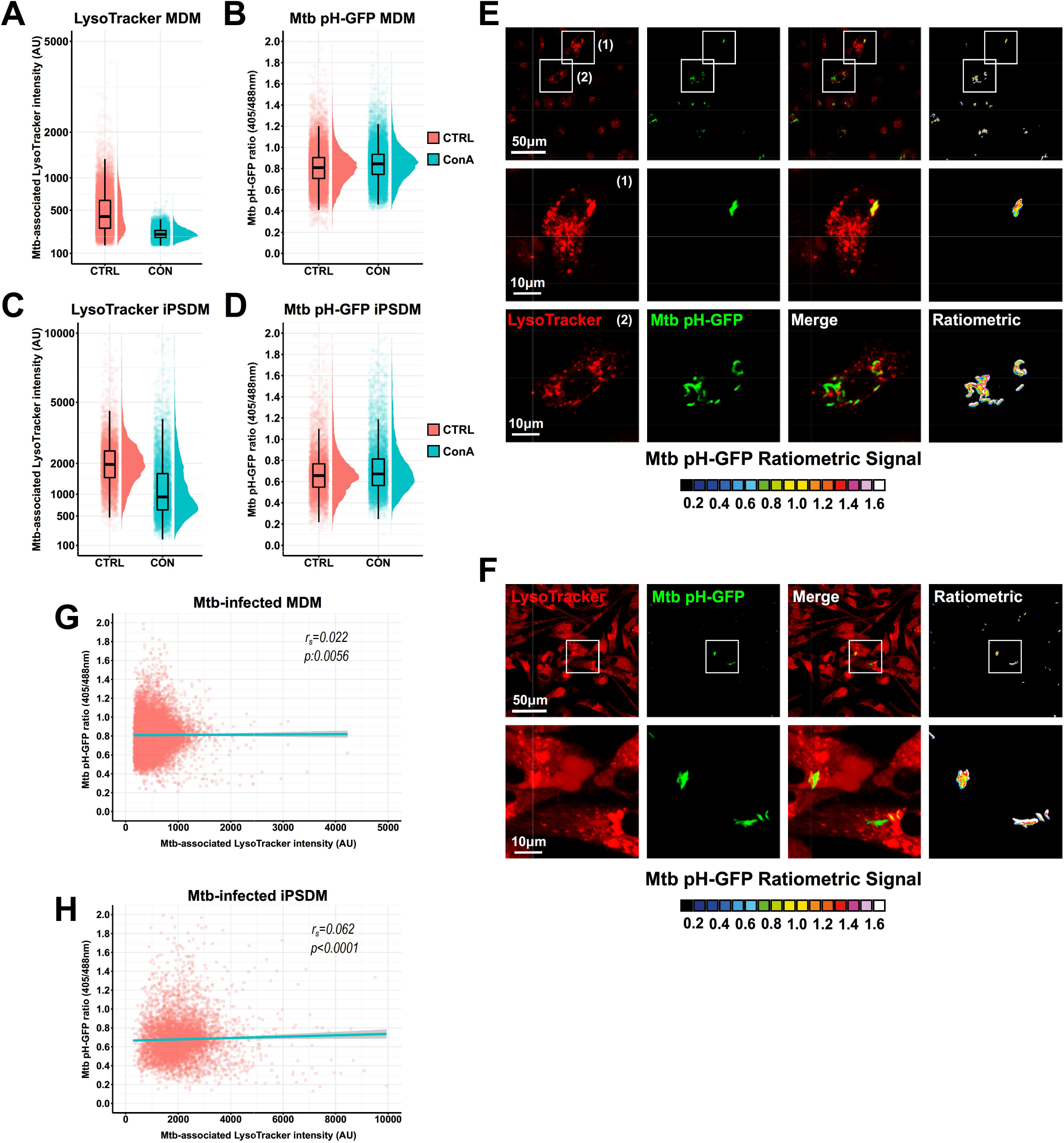
High content dual-imaging of Mtb intrabacterial pH and host-cell intracellular acidification. Human macrophages were infected for 24 hours and subsequently treated with ConA or left untreated for an additional 24 hours. Cells were then pulsed with 200 nM of LysoTracker Red for 30 min before live-acquisition was performed using the OPERA Phenix imaging platform. Quantitative analysis of Mtb pH-GFP ratio (405/488nm) and Mtb-associated LysoTracker were performed using the Harmony software. **(A-C)** Quantification of Mtb-associated LysoTracker mean fluorescence intensities within **(A)** infected-MDM or **(C)** infected-iPSDM in the absence (pink) or presence (cyan) of v-ATPase inhibitor ConA. Results obtained from CTRL or ConA-treated samples are displayed as raincloud plots where black box-plots are overlaid on top of individual raw data and associated with their respective density plots. **(B-D)** Quantification of Mtb pH-GFP ratio (405/488nm) within **(B)** infected-MDM or **(D)** infected-iPSDM in the absence (pink) or presence (cyan) of v-ATPase inhibitor ConA. Results obtained from CTRL or ConA-treated samples are displayed as raincloud plots where black box-plots are overlaid on top of individual raw data and associated with their respective density plots. **(E-F)** Representative micrographs display LysoTracker labelling (red) and Mtb pH-GFP (green). Ratiometric signal was obtained by dividing the fluorescence intensity acquired with excitation/emission channels of 405/510 nm by the one obtained at 488/510 nm. Ratiometric signal is displayed as a 16-colour palette ranging from 0 to 1.6 units. Scale bar corresponds to 50 µm. Panel **(E)** shows MDM and panel **(F)** iPSDM. Regions of interest highlighted by the white rectangles, are shown in detail in the bottom panels respectively. Scale bar corresponds to 10 µm. **(G-H)** Spearman’s correlation between Mtb-associated LysoTracker (x-axis) and Mtb pH-GFP ratio (405/488nm) (y-axis) signals in individual bacterial region of interests within **(G)** infected-MDM or **(H)** infected-iPSDM. The cyan line shows the linear regression model, the Spearman rank correlation coefficient (*r*_*s*_) and the corresponding *p*-value were calculated by using the ggpubr R package and two-tailed statistical t-test. Between 5895 and 15823 bacterial regions of interest were analysed per experimental condition. Results are representative are from n = 2 biologically independent experiments performed at least in two-three technical replicates.

### Mtb subcellular localisation within acidic compartments and time of residence does not affect bacterial pH homeostasis

To complement our live-snapshot imaging approach and capture the dynamic nature of these transient events, we performed live imaging at low-content/high resolution and tracked individual mycobacterial regions of interest (mROI) to define whether the time of residency within LysoTracker positive compartments impacts Mtb intrabacterial pH. Live cell imaging of Mtb-infected iPSDM was performed over a 6 h period with 30 min intervals to minimise photobleaching and/or phototoxicity. Monitoring and tracking of mROI revealed at least 4 different phenotypic profiles: (i) LysoTracker^(-)^ Mtb that became LysoTracker^(+)^ (**Figure 2A**), (ii) LysoTracker^(+)^ Mtb that remained LysoTracker^(+)^ (**Figure 2B**), (iii) LysoTracker^(+)^ Mtb that became LysoTracker^(-)^ (**Figure 2C**), and (iv) LysoTracker^(-)^ Mtb that remained LysoTracker^(-)^ (**Figure 2D**). Next, we analysed the dynamics of Mtb-associated LysoTracker intensity and pH-GFP ratio (**Figure 2E**) as previously described (Bernard et al., 2020; Schnettger et al., 2017). We observed that over time the association with LysoTracker was very heterogenous and that Mtb pH-GFP ratio did not significantly change, with no correlation between LysoTracker association and Mtb pH-GFP ratio (**Figure 2E**). To exclude that prolonged exposure to low pH within an acidified compartment impacts Mtb pH homeostasis, we analysed the cumulative values of Mtb-associated LysoTracker fluorescence intensity over time to include total LysoTracker intensity association during 360 min and the corresponding Mtb pH-GFP ratios (**Figure 2F**). Thus, giving a quantitative profile of LysoTracker intensity faced by multiple mROI over the entire time-course. In agreement with the previous analysis, despite heterogenous accumulation of LysoTracker with Mtb, there was no significant changes in Mtb pH homeostasis (**Figure 2F**). Determination of Spearman’s correlation coefficients at the end time point, did not show positive or negative association between subcellular acidification profile and Mtb intrabacterial pH with a value of *r*_*s*_ = -0.075 and *r*_*s*_ = -0.011 for the two different analyses respectively (**Figure 2E and 2F**). We concluded that Mtb is able to maintain its intracellular pH even when facing fluctuating acidic intracellular environments (Darby et al., 2013; Fontes et al., 2020; Vandal et al., 2008).

**Figure 2:**
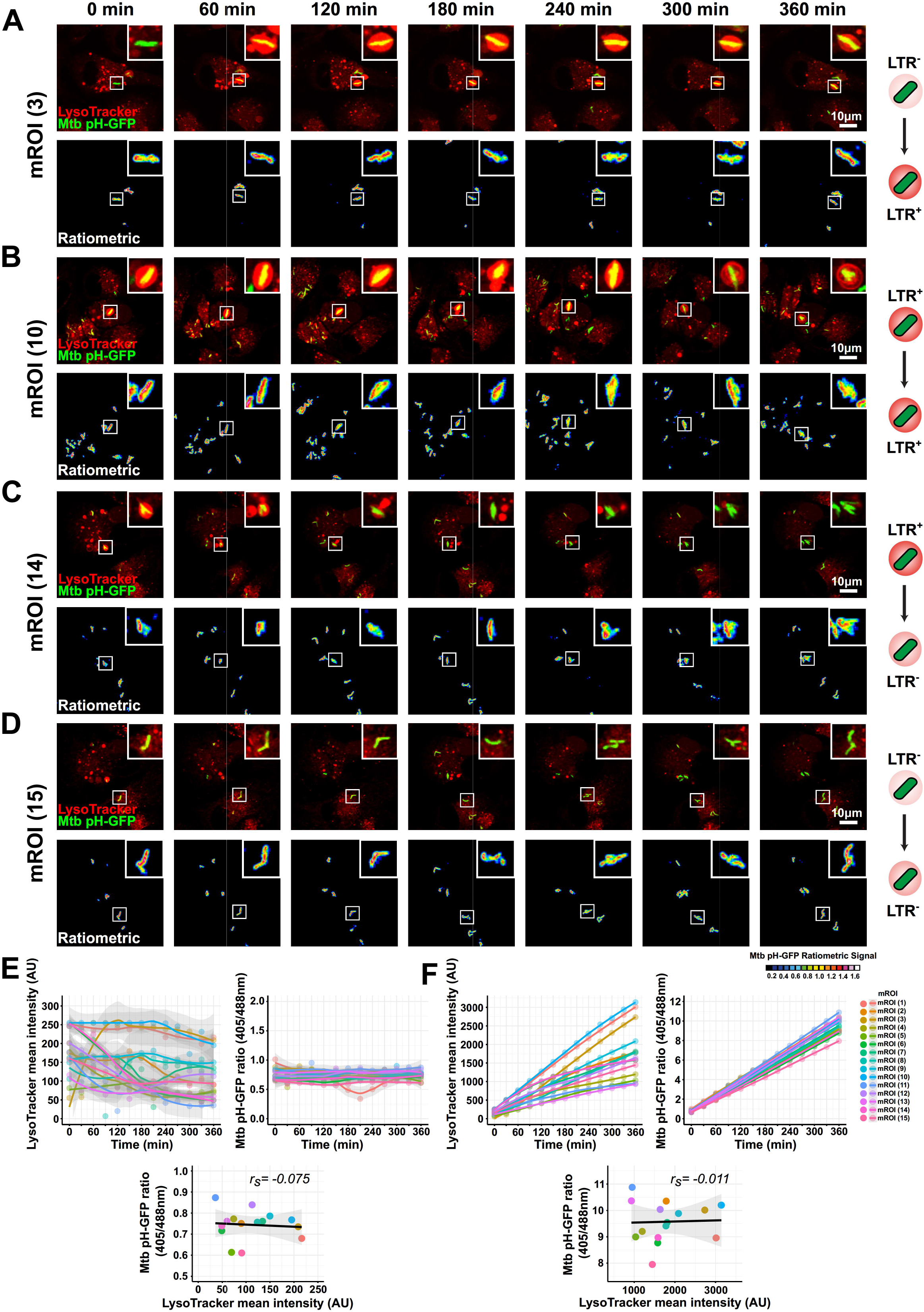
Live-cell imaging of Mtb pH homeostasis in acidic subcellular compartments. Human iPSDM were infected for 24 hours and then pulsed with 200 nM of LysoTracker Red for 30 min before live-acquisition was performed using a Leica SP5 AOBS Laser Scanning Confocal Microscope. Quantitative analysis of Mtb pH-GFP ratio (405/488nm) and Mtb-associated LysoTracker were performed using the open source Fiji software. **(A-D)** Representative micrographs display LysoTracker labelling (red) and Mtb pH-GFP (green) of 4 distinct mROI (3, 10, 14 and 15 respectively) with different LysoTracker associated fluorescence patterns along the kinetic. Ratiometric signal was obtained by dividing the fluorescence intensity acquired with excitation/emission channels of 405/510 nm by the one obtained at 488/510 nm. Ratiometric signal is displayed as a 16-colour palette ranging from 0 to 1.6 units. Events of interest are highlighted with white squares and a zoom in is displayed at the top right corner of each micrograph. Scale bar corresponds to 10 µm. **(E)** Quantitative analysis of intracellular Mtb-associated LysoTracker intensity and Mtb pH-GFP profiles of single-tracked mROI over time. Left panel shows the mean LysoTracker intensity associated to Mtb pH-GFP and the right panel shows their corresponding fluorescence ratio profiles. Bottom panel shows Spearman’s correlation between Mtb-associated LysoTracker (x-axis) and Mtb pH-GFP ratio (405/488nm) (y-axis) signals from single-tracked mROI at the end of the kinetic (t_360 min_). Spearman rank correlation coefficient (*r*_*s*_) was calculated by using the ggpubr R package. Each colour represents one mROI and the corresponding legend is displayed in **(F). (F)** Quantitative analysis of cumulative Mtb-associated LysoTracker intensity and Mtb pH-GFP profiles of single-tracked mROI over time. Left panel shows the cumulative mean LysoTracker intensity associated to Mtb pH-GFP and the right panel shows their corresponding fluorescence ratio profiles. Bottom panel shows Spearman’s correlation between cumulative Mtb-associated LysoTracker values (x-axis) and cumulative Mtb pH-GFP ratio values (405/488nm) (y-axis) signals from single-tracked mROI over the kinetic (t_360 min_). Spearman rank correlation coefficient (*r*_*s*_) was calculated by using the ggpubr R package. Each colour represents one mROI and the corresponding legend is displayed on the right of the panel. Results are from n = 15 individually tracked mROI.

### Spatiotemporal analysis of PZA-mediated Mtb intrabacterial pH homeostasis disruption *in cellulo*

Experimental investigations of the molecular mechanism(s) of PZA action have been mostly performed within *in vitro* cell-free media and it is unknown whether PZA/POA molecules act as bacterial protonophores to disrupt Mtb intrabacterial pH homeostasis *in cellulo*. To address this question, human macrophages were infected with Mtb pH-GFP for 24 h and subsequently treated with PZA, BDQ, INH, RIF or left untreated for an additional 24 h before being stained with LysoTracker and live-imaged. Quantitative analysis of Mtb-associated LysoTracker intensity revealed that BDQ treatment was the only condition impacting Mtb-associated LysoTracker intensity profile as previously reported (median_BDQ_ = 4252.1; IQR_BDQ_ = 3369.3 and median_CTRL_ = 2502.2; IQR_CTRL_ = 1221.3, respectively) (**Figure 3A**) (Giraud-Gatineau et al., 2020). However, despite its proposed ionophore activity *in vitro* (Hards et al., 2018) and potent activity of enhancing the intracellular acidification processes (Giraud-Gatineau et al., 2020), such effects were not sufficient to overcome bacterial regulation of cytosolic pH, as we did not observe changes in Mtb intrabacterial pH in the presence of BDQ (mean normalised pH-GFP ratio of -0.006; *p*-value = 0.887) (**Figure 3B-C**). Similar results were obtained in the presence of INH and RIF where Mtb pH-GFP ratios were similar to the untreated control condition (mean normalised pH-GFP ratios of 0.016; *p*-value = 0.160 and -0.014; *p*-value = 0.319, respectively) (**Figure 3B-C**). Strikingly, from the four different antibiotics tested, PZA was the only one able to induce changes in Mtb pH-GFP ratio, providing evidence that PZA displays intrabacterial pH-disruptive activity in Mtb-infected human macrophages (mean normalised pH-GFP ratio of 0.155; *p*-value ≤0.001) (**Figure 3C**). PZA/POA molecules require endolysosomal acidification to accumulate inside Mtb and display antimicrobial efficacy (Santucci et al., 2021). We hypothesized that this process is likely resulting from the conversion of POA^(-)^ into its protonated form HPOA within acidic host-microenvironments (**Figure 3D**). To investigate whether PZA/POA-mediated intrabacterial pH homeostasis disruption *in cellulo* requires endolysosomal acidification, Mtb-infected MDM were treated with increasing concentration of PZA ranging from 0 to 400 mg/L in the presence or absence of the v-ATPase inhibitor ConA and both host and bacterial acidification profiles were monitored at 4 h, 16 h, 24 h and 72 hours post-treatment (**Figure 3E**) using our live dual-imaging approach. Quantitative analysis of Mtb pH-GFP fluorescence profiles at the indicated time points in untreated control cells confirmed that Mtb can stably maintain intrabacterial pH through the course of the infection (**Figure 3F-I**). Treatment of infected human macrophages with PZA was able to decrease Mtb intrabacterial pH in a time-and concentration-dependent manner (**Figure 3F-I**). After 4 h of treatment, only 400 mg/L of PZA showed a detectable effect on Mtb intrabacterial pH (mean normalised pH-GFP ratio of 0.0625; *p*-value ≤ 0.01) (**Figure 3F**). After 16 h, 24 h and 72 h, PZA concentrations ranging from 30 mg/L to 400 mg/L significantly disrupted bacterial pH homeostasis (**Figure 3G-3I**). The absolute changes in Mtb pH-GFP ratio relative to the control condition, confirmed a time- and concentration-dependent effect of PZA on Mtb intrabacterial pH (**Figure 3J-M**). Importantly, PZA treatment did not induce rerouting of Mtb into endolysosomal compartments (**Figure S3**), ruling out a concentration-dependent effect towards pH-GFP ratio due to excessive lysosomal delivery. We also noticed that the LysoTracker intensity profile decreased overtime independently of the infection, suggesting that human primary macrophages display optimal lysosomal activity for a limited amount of time during their *in vitro* lifespan (**Figure S4**). ConA co-treatment with increasing concentrations of PZA resulted in an almost complete loss of PZA Mtb pH-disruptive function (**Figure 3J-M**) suggesting that functional acidification of the Mtb phagosome is a prerequisite for PZA-mediated pH disruption in intracellular Mtb. Quantitative correlative analysis of Mtb pH-GFP ratio values with associated LysoTracker intensity at increasing PZA concentrations did not show a direct association, with Spearman’s correlation coefficients *r*_*s*_ between - 0.3 and 0.3 (**Figure S5**). These data highlight that PZA-mediated pH homeostasis disruption within acidic environments is a dynamic process.

**Figure 3:**
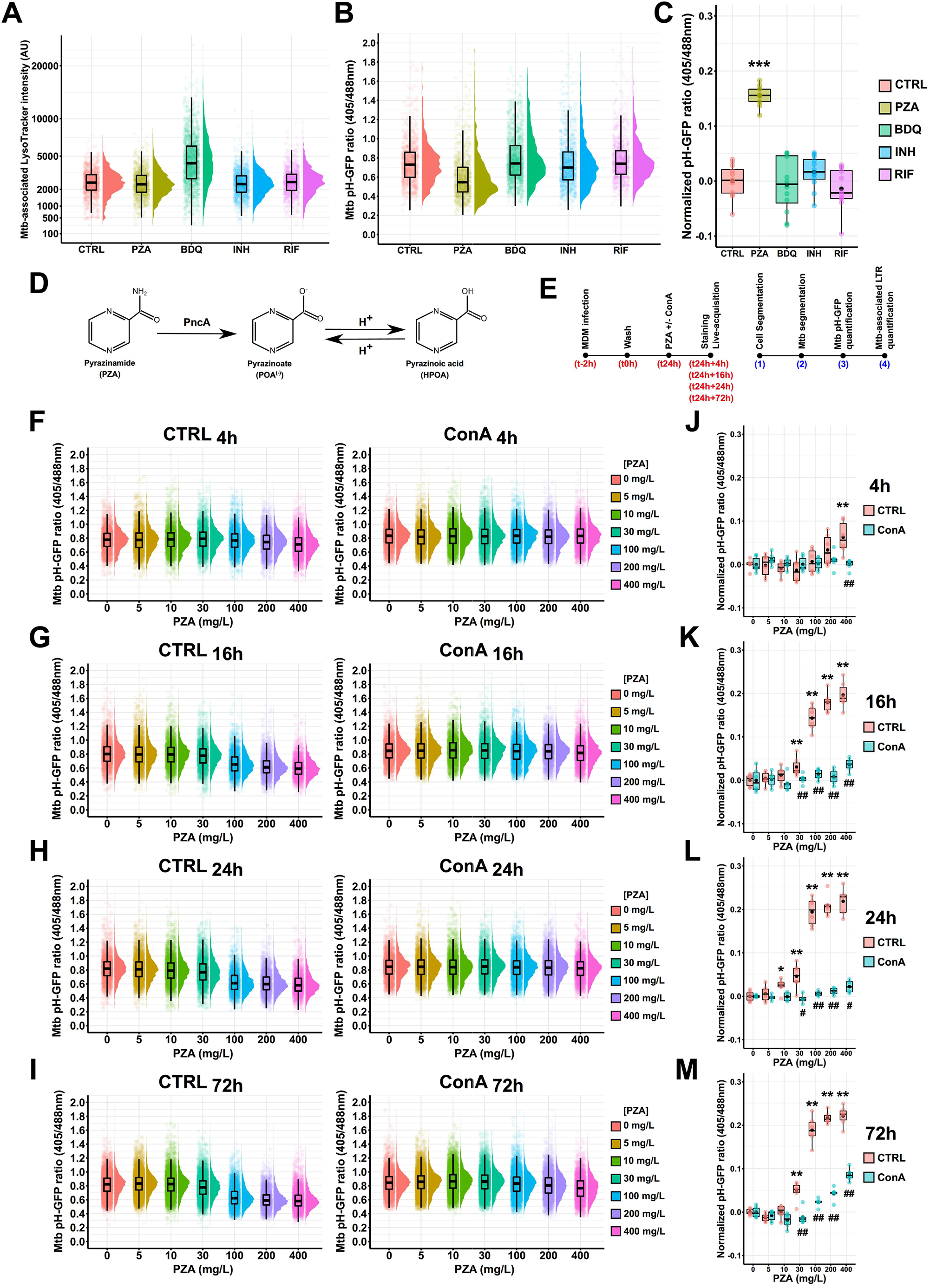
Spatiotemporal analysis of PZA-mediated Mtb intrabacterial pH homeostasis disruption *in cellulo*. **(A-C)** Human macrophages were infected with Mtb pH-GFP for 24 hours and subsequently treated with either 100 mg/L of PZA, 2.5 mg/L of BDQ, 5 mg/L of INH, 5 mg/L of RIF or left untreated for 24 hours. Cells were then pulsed with 200 nM of LysoTracker Red for 30 min before live-acquisition was performed using the OPERA Phenix imaging platform. **(A)** Quantification of Mtb-associated Lysotracker mean intensity or **(B)** Mtb pH-GFP ratio (405/488nm) within infected-MDM treated with different antibiotics for 24 hours. Results are displayed as raincloud plots where black box-plots are overlaid on top of individual raw data and associated with their respective density plots. Between 1102 and 1658 bacterial regions of interest were analysed per experimental condition. **(C)** Determination of absolute changes in Mtb pH-GFP ratio (405/488nm) upon various antibiotic treatment. Mean pH-GFP ratio of each antibiotic treatment was subtracted from the untreated control 24-hours post-treatment to obtain an absolute value reflecting antibiotic-mediated pH disruption normalized to the control. Determination of normalized pH-GFP ratio was performed for each condition and results are displayed as box-plots with individual replicate data. Black dots were added to highlight the mean of each conditions. Each colour represents a specific antibiotic or the control condition. Results are from n = 2 biologically independent experiments performed at least in two-three technical replicates. Statistical analysis was performed using Wilcoxon signed-rank test where untreated control was used as reference condition (where ^*^ *p* ≤ 0.05; ^**^ *p* ≤ 0.01; ^***^ *p* ≤ 0.001). **(D)** Chemical structures of PZA, POA^(-)^ and HPOA. Conversion of the prodrug PZA into POA^(-)^ is mediated by the bacterial pyrazinamidase PncA and transition of POA^(-)^ into HPOA is driven by proton availability. **(E)** Schematic representation of the experimental procedure followed to perform MDM infection, staining and fluorescence microscopy imaging. **(F-M)** Human macrophages were infected for 24 hours and subsequently treated with increasing concentration of PZA ranging from 0-400 mg/L in absence or presence of ConA for 4 h, 16 h, 24 h or 72 hours. Cells were then pulsed with 200 nM of LysoTracker Red for 30 min before live-acquisition was performed using the OPERA Phenix imaging platform. **(F-I)** Quantification of Mtb pH-GFP ratio (405/488nm) within infected-MDM treated with increasing concentration of PZA ranging from 0-400 mg/L in the absence or presence of v-ATPase inhibitor ConA at 4 h, 16 h, 24 h or 72 hours post-treatment. Results are displayed as raincloud plots where black box-plots are overlaid on top of individual raw data and associated with their respective density plots. Each colour represents a specific PZA concentration. Between 2164 and 8813 bacterial regions of interest were analysed per experimental condition **(J-M)** Determination of absolute changes in Mtb pH-GFP ratio (405/488nm) upon PZA treatment. Mean pH-GFP ratio of each antibiotic treatment was subtracted from the PZA-untreated control to obtain an absolute value reflecting antibiotic-mediated pH disruption normalized to its respective control in the presence or absence of ConA at 4 h, 16 h, 24 h or 72 hours post-treatment. Determination of normalized pH-GFP ratio was performed in the absence (pink) or presence (cyan) of v-ATPase inhibitor ConA. Results are displayed as box-plots with individual data. Black dots were added to highlight the mean of each conditions. Results are from n = 2 biologically independent experiments performed at least in two-three technical replicates. Statistical analysis was performed using Wilcoxon signed-rank test where PZA effect on Mtb intrabacterial pH was assessed against the untreated control (where ^*^ *p* ≤ 0.05; ^**^ *p* ≤ 0.01; ^***^ *p* ≤ 0.001) and ConA effect towards PZA was assessed by comparing each concentration with its respective untreated control as reference condition (where ^#^ *p* ≤ 0.05; ^##^ *p* ≤ 0.01; ^*###*^*p* ≤ 0.001).

### Endolysosomal acidification and protonophore activity of PZA contribute to Mtb restriction in human macrophages

We next hypothesized that functional host intracellular acidification and PZA-mediated pH-decrease are required for mycobacterial growth inhibition. In order to test this hypothesis, we quantified Mtb intracellular replication at the single-cell level in the presence of increasing concentration of PZA in both control or ConA treated cells (**Figure 4A**). Results from dose-response analysis showed that functional endolysosomal acidification, required for PZA-mediated pH disruption, is also required for optimal PZA efficacy and Mtb growth restriction over the course of infection (**Figure 4A**). The determination of PZA half maximal effective concentration (EC_50_) using a four-parameter logistic non-linear regression model showed that ConA co-treatment increased, by approximately 3.5 times, the amount of antibiotic required to efficiently inhibit 50% of Mtb growth *in cellulo* (49.5 ± 19.2 mg/L and 173.1 ± 35.2 mg/L, respectively) (**Figure 4A-4B**). These results agree with our previous observations showing that the use of v-ATPase inhibitors is able to counteract PZA/POA-mediated growth inhibition by impairing POA accumulation within the bacteria (Santucci et al., 2021). These experiments were also performed in another human macrophage model using iPSDM (**Figure S6**). Notably, in iPSDM antagonistic effects between PZA and ConA were also observed, however the phenotypes were less pronounced than in Mtb-infected MDM suggesting that iPSDM and MDM might have different intracellular pH homeostatic processes. Altogether, these findings support the proposed pH-dependent mode of action of PZA, in which endolysosomal acidification is a necessary prerequisite and driver of the protonophore activity of PZA/POA, which controls bacterial growth in human macrophages.

**Figure 4:**
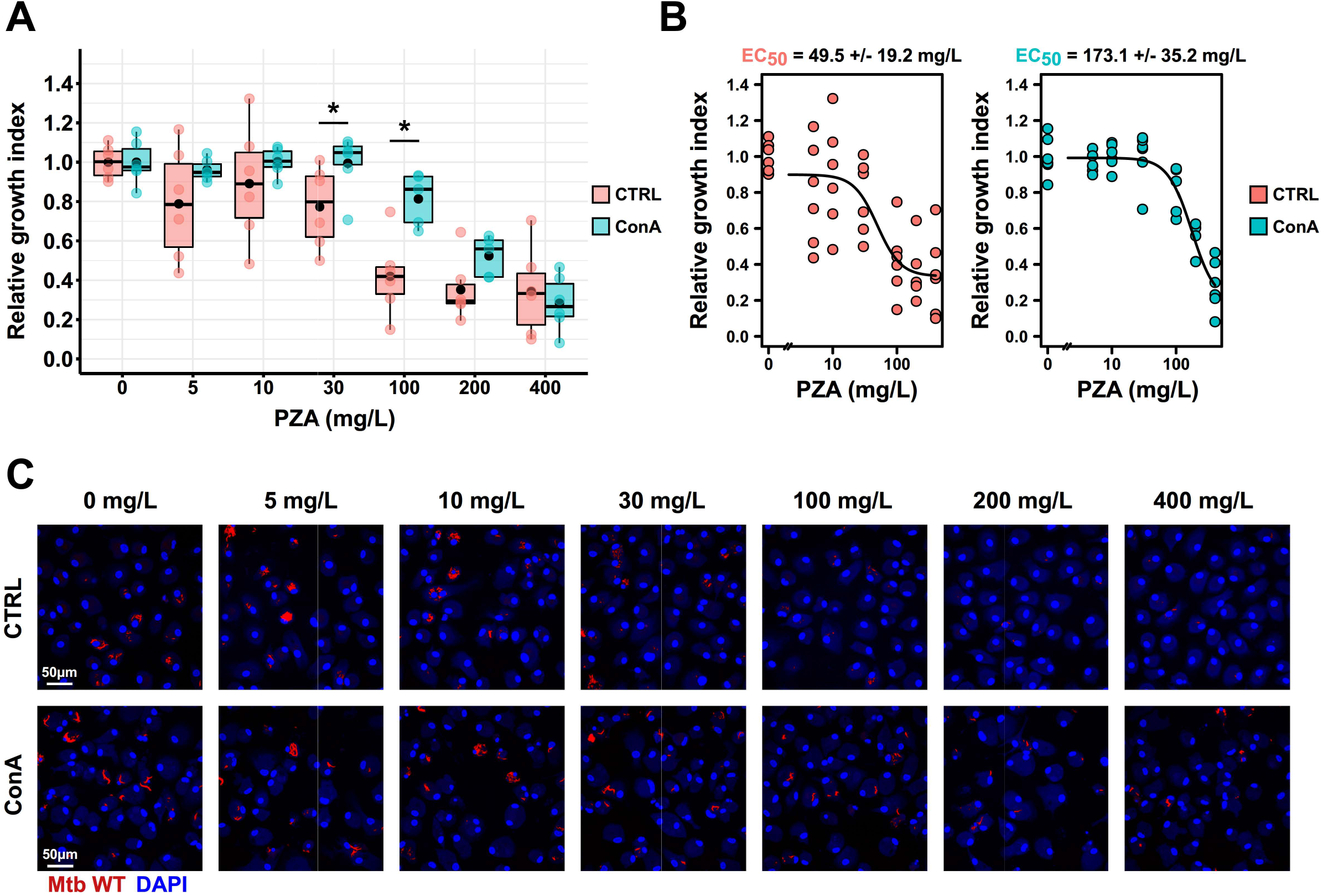
PZA-mediated growth inhibition requires endolysosomal acidification within human macrophages. Human macrophages were infected with Mtb WT E2-Crimson for 24 hours and subsequently treated with increasing concentration of PZA ranging from 0-400 mg/L in the absence or presence of ConA for 72 hours. **(A)** Quantitative analysis of E2-Crimson Mtb WT replication at the single cell level within MDM treated with increasing concentration of PZA in the absence or presence of ConA. Normalization was done to the mean Mtb area per cell pre-treatment (t_24h_ post-infection) and the control condition without PZA was used as reference corresponding to 100 % growth. Determination of relative growth index was performed in the absence (pink) or presence (cyan) of v-ATPase inhibitor ConA. Results are displayed as box-plots with individual replicate data. Black dots were added to highlight the mean of each condition. Between 1185 and 1925 Mtb-infected MDM were analysed by high-content single-cell microscopy and results are representative from n = 2 biologically independent experiments performed at least in two-three technical replicates. Statistical analysis was performed using Wilcoxon signed-rank test where untreated control was used as reference condition (where ^*^*p* ^*^ 0.05; ^**^ *p* ≤ 0.01; ^***^ *p* ≤ 0.001) **(B)** Determination of PZA EC_50_ in the absence or presence of ConA by performing a 4-parameter nonlinear logistic regression of the data displayed in **(A). (C)** Representative confocal fluorescence images of Mtb WT-infected MDM for 24 hours and further treated for 72 hours with increasing concentrations of PZA. Magnifications display nuclear staining (blue) and Mtb-producing E2-Crimson (red). Scale bar corresponds to 50 µm. Micrographs are representative of 2 independent experiments.

### Mtb mutants with different subcellular localisations show distinct PZA susceptibility profiles *in cellulo*

To define how intracellular localisation contributes to PZA/POA antibacterial efficacy, we assessed PZA-mediated pH homeostasis disruption and growth inhibition towards multiple Mtb mutants with distinct intracellular lifestyles (**Figure 5A**). Mtb WT harbouring a functional ESX-1 secretion system was used as the reference strain and Mtb ΔRD1 lacking a functional ESX-1 machinery was used as a phagosome-restricted strain. We also included another phagosome-restricted mutant, Mtb Δ*esxBA* which lacks only the two major ESX-1 effectors EsxA and EsxB (also known as ESAT-6 and CFP-10). A relative growth index was quantified for each strain, and a dose-response analysis using a four-parameter logistic non-linear regression model was performed to determine PZA EC_50_ towards each strain (**Figure 5B-5D**). Results obtained after curve fitting showed that EC_50_ towards the reference strain Mtb WT was 33.8 ± 8.5mg/L. Both Mtb ΔRD1 and Mtb Δ*esxBA* displayed increased susceptibility to PZA with EC_50_ values of 13.0 ± 3.7 and 17.8 ± 7.2 mg/L respectively. Determination of EC_50_ values further highlighted the increased in susceptibility of strains unable to damage the endolysosomal membrane and access host cytosol with a 2.60-and 1.90-fold increase for Mtb ΔRD1 and Mtb Δ*esxBA* respectively, when compared to the WT reference strain (**Figure 5C**). Thus, a functional ESX-1 secretion system is protective against PZA/POA activity in Mtb infected macrophages, potentially through facilitating membrane damage and cytosolic access. Previous work showed that disruption of Mtb intrabacterial pH homeostasis caused by pharmacological inhibitors directly correlated with a mycobactericidal effect (Darby et al., 2013). We investigated whether a PZA-mediated pH decrease was correlated with intracellular growth defects. Relative growth index values were plotted as a function of normalised pH-GFP ratios at each PZA concentration and Spearman’s correlation coefficients were determined for each strain (**Figure 5E**). Notably, PZA-mediated pH homeostasis disruption strongly correlated with the intracellular replication defect (*r*_*s*_ values ranging from -0.82 to -0.86) suggesting that PZA-mediated intrabacterial pH disruption is an important factor in its antibacterial activity (**Figure 5E**).

**Figure 5:**
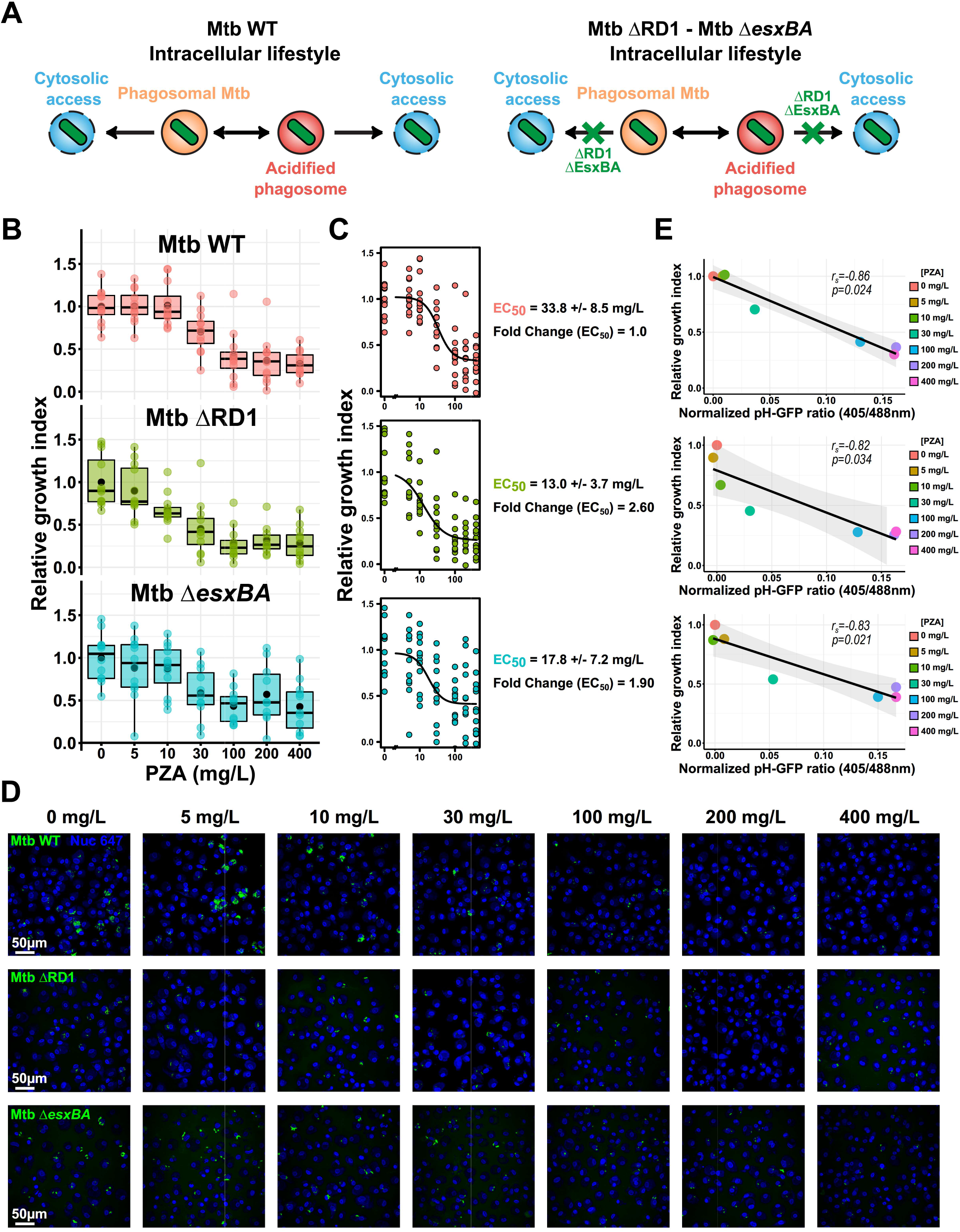
RD1-and EsxBA-mediated endolysosomal damage partially protects Mtb against PZA activity *in cellulo*. Human macrophages were infected with Mtb WT, Mtb ΔRD1 or Mtb Δ*esxBA* for 24 hours and subsequently treated with increasing concentrations of PZA ranging from 0-400 mg/L for 72 hours. **(A)** A schematic representation of Mtb WT, Mtb ΔRD1 and Mtb Δ*esxBA* intracellular lifestyles. The contribution of RD1 and EsxBA virulence factors in membrane damage and cytosolic access is highlighted in the right panel. **(B)** Quantitative analysis of fluorescent Mtb strain replication at the single cell level within MDM treated with increasing concentrations of PZA. Normalization was done to the mean Mtb area per cell pre-treatment (t_24h_ post-infection) and the control condition without PZA was used as reference corresponding to 100 % growth. Results are displayed as box-plots with individuals replicate data. Black dots were added to highlight the mean of each conditions. Between 3880 and 10874 Mtb-infected MDM were analysed by high-content single-cell microscopy and results are representative from n = 4 biologically independent experiments performed at least in two-three technical replicates. **(C)** Determination of PZA EC_50_ for the different Mtb strains by performing a 4-parameter nonlinear logistic regression of the data displayed in **(B). (D)** Representative confocal fluorescence images of Mtb WT, Mtb ΔRD1 or Mtb Δ*esxBA*-infected MDM for 24 hours and further treated for 72 hours with increasing concentrations of PZA. Magnifications display nuclear staining (blue) and fluorescent Mtb (green). Scale bar corresponds to 50 µm. Micrographs are representative of 4 independent experiments. **(E)** Spearman’s correlation between normalized Mtb pH-GFP ratio (405/488nm) at 24 hours post-treatment (x-axis) and Mtb relative growth index (y-axis) within infected-MDM at 72 hours post-treatment. Results from Mtb WT, Mtb ΔRD1 and Mtb Δ*esxBA* are shown from top panel to the bottom panel respectively. The black line shows the linear regression model, the Spearman rank correlation coefficient (*r*_*s*_) and the corresponding *p*-value were calculated by using the ggpubr R package and two-tailed statistical t-test. Results are from n = 2 or n = 4 biologically independent experiments performed at least in two-three technical replicates.

### PZA-mediated intrabacterial pH disruption and growth inhibition *in cellulo* requires POA conversion by functional PncA

Finally, we sought to understand whether PZA conversion to POA was essential to display its pH-disruptive property and antibacterial capacity. To answer this question, we used the bovine TB agent and zoonotic pathogen, *Mycobacterium bovis* (Mbv). As a member of the *Mycobacterium tuberculosis* complex, Mbv was chosen due to its ability to replicate within human macrophages (Queval et al., 2021) and its well-characterised intrinsic resistance towards PZA (Petrella et al., 2011; Scorpio and Zhang, 1996). Indeed, Mbv harbours a point mutation within its *pncA* gene that is responsible for the H57D substitution, which blocks PZA to POA conversion (Petrella et al., 2011; Scorpio and Zhang, 1996). We generated a pH-GFP reporter Mbv strain (Mbv pH-GFP) and assessed whether PZA-mediated pH disruption was occurring in Mbv-infected MDM. As expected, without PZA to POA conversion, no protonophore activity was noticeable against Mbv (**Figure 6A and 6D**) even at 400mg/L. In addition, PZA intracellular activity on Mbv replication was also investigated by quantitative fluorescence microscopy. In agreement with previous reports from *in vitro* studies (Scorpio and Zhang, 1996), Mbv was resistant to PZA *in cellulo* (**Figure 6B, 6C and 6E**). These results demonstrate that *in cellulo*, the protonophore activity of PZA is mediated by its active form POA and that such conversion is essential for antibacterial activity within infected macrophages.

**Figure 6:**
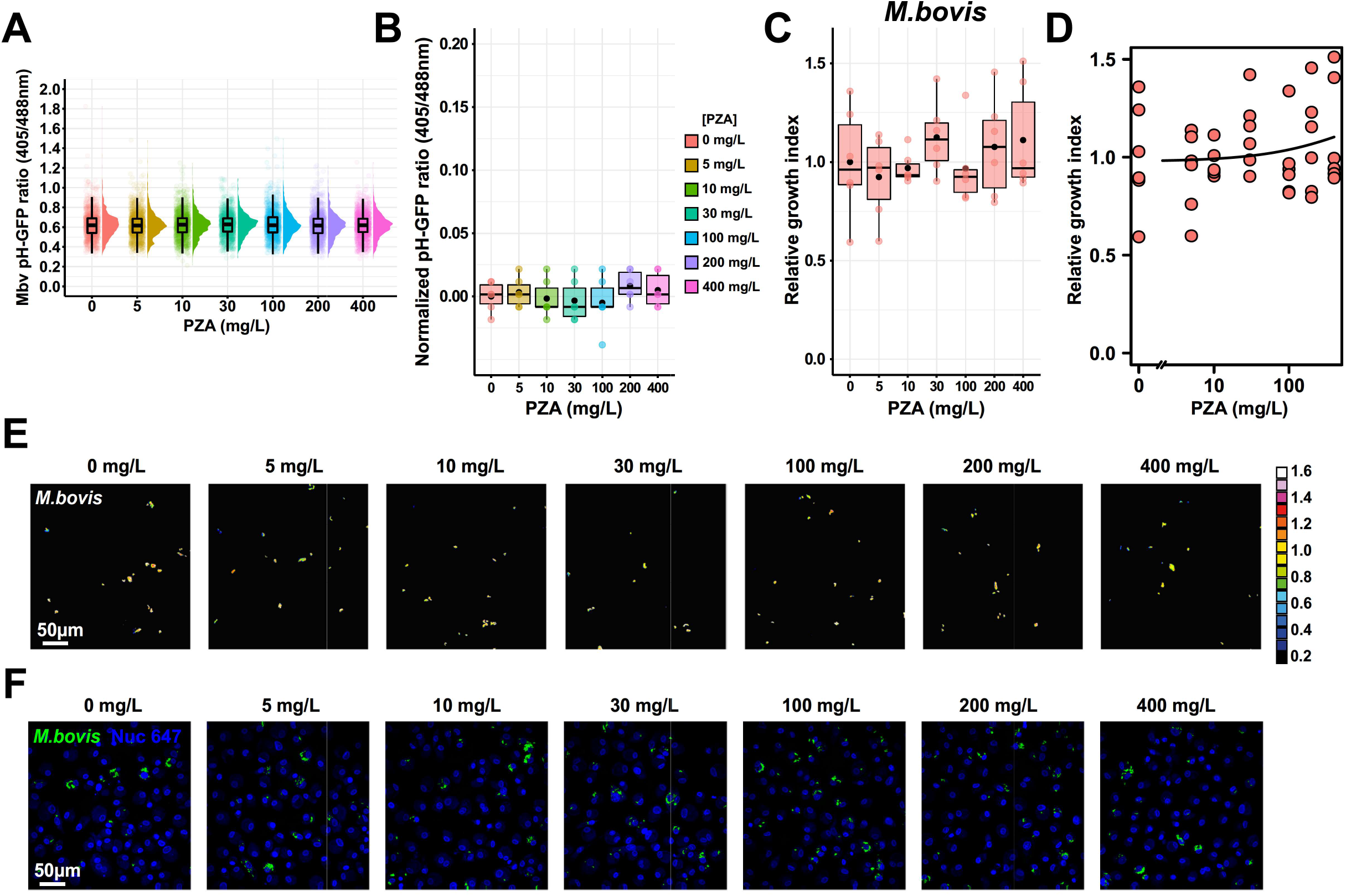
PZA-mediated intrabacterial pH homeostasis disruption and growth inhibition *in cellulo* requires functional PncA. Human macrophages were infected with Mbv pH-GFP, for 24 hours and subsequently treated with increasing concentrations of PZA ranging from 0-400 mg/L for 24 hours. Cells were then pulsed with 200 nM of LysoTracker Red for 30 min before live-acquisition was performed using the OPERA Phenix imaging platform. **(A)** Quantification of Mbv pH-GFP ratio (405/488nm) within infected-MDM treated with increasing concentration of PZA ranging from 0-400 mg/L at 24 hours post-treatment. Results are displayed as raincloud plots where black box-plots are overlaid on top of individual replicate data and associated with their respective density plots. Each colour represents a specific PZA concentration. **(B)** Determination of absolute changes in Mbv pH-GFP ratio (405/488nm) from upon PZA treatment. Determination of absolute changes in Mtb pH-GFP ratio (405/488nm) upon various antibiotic treatment. Mean pH-GFP ratio of each antibiotic treatment was subtracted from the untreated control 24-hours post-treatment to obtain an absolute value reflecting antibiotic-mediated pH disruption normalized to the control. Results are displayed as box-plots with individuals replicate data. Black dots were added to highlight the mean of each condition. **(C-D)** Human macrophages were infected with fluorescent Mbv for 24 hours and subsequently treated with increasing concentrations of PZA ranging from 0-400 mg/L for 72 hours. **(C)** Quantitative analysis of fluorescent Mtb strains replication at the single cell level within MDM treated with increasing concentrations of PZA. Normalization was done to the mean Mbv area per cell pre-treatment (t_24h_ post-infection) and the control condition without PZA was used as reference corresponding to 100 % growth. Results are displayed as box-plots with individuals’ raw data. Black dots were added to highlight the mean of each condition. Between 4072 and 4628 Mbv-infected MDM were analysed by high-content single-cell microscopy and results are representative from n = 2 biologically independent experiments performed at least in two-three technical replicates. **(D)** Determination of PZA EC_50_ for the Mbv strain by performing a 4-parameter nonlinear logistic regression of the data displayed in **(C). (E)** Representative fluorescence ratiometric images of Mbv-infected MDM infected for 24 hours and further treated for 24 hours with increasing concentrations of PZA. Ratiometric signal is displayed as a 16-colour palette ranging from 0 to 1.6 units. Scale bar corresponds to 50 µm. Micrographs are representative of 2 independent experiments. **(F)** Representative confocal fluorescence images of Mbv-infected MDM for 24 hours and further treated for 72 hours with increasing concentrations of PZA. Magnifications display nuclear staining (blue) and fluorescent Mbv (green). Scale bar corresponds to 50 µm. Micrographs are representative of 2 independent experiments.

## Discussion

Here we described a novel live dual imaging approach to monitor pH homeostasis within both the host cell and the pathogen in a biosafety level 3 laboratory. This approach is applicable to both human primary monocyte derived macrophages (MDM) and human iPS-derived macrophages (iPSDM). In both MDM and iPSDM, ConA treatment was able to reduce Mtb-associated LysoTracker intensity suggesting that this pharmacological inhibition is a powerful approach to perturb endolysosomal acidification in these two macrophage models. Quantitative analysis revealed that the regulation of Mtb intrabacterial pH *in cellulo* is not homogeneous with a subset of bacilli displaying a differential pH regulation as shown in *E. coli in vitro* (Goode et al., 2021). At the intra-macrophage population level, our data show that Mtb can maintain its own pH within endolysosomes of human macrophages, in agreement with previous findings *in vitro* and in mouse macrophages treated or not with interferon-γ (Darby et al., 2013; Fontes et al., 2020; Vandal et al., 2008). An important finding of our analysis is that impairment of endolysosomal acidification had no impact on intracellular Mtb pH homeostasis. Despite a very significant heterogeneity in LysoTracker association, most bacteria maintained their intracellular pH during the course of infection, suggesting that Mtb is very efficient at regulating intrabacterial pH in its preferred host cell.

Intracellular Mtb continuously switches between membrane bound (able to retain protons and acidic) and cytosolic (host-cell neutral pH) localisation (Bussi and Gutierrez, 2019) and here by using live imaging of subcellular acidification, we tested the hypothesis that when residing in an acidic environment, the exposure to an acidic pH will reduce intrabacterial pH. Unexpectedly, we found that irrespective of the time of residence in an acidic compartment, the bacilli maintained a constant intrabacterial pH, suggesting that mycobacteria have potent mechanisms of pH sensing and homeostasis in fluctuating environments and can rapidly respond to changes (Krulwich et al., 2011; Vandal et al., 2009). A limitation of our study is that we are not able to define if these bacteria in an acidic host compartment are alive and able to replicate, in a non-replicating state or eventually dead. More studies and technical developments are required to explore this in detail. It is also important to mention that LysoTracker staining doesn’t allow to clearly discriminate between pH 4.5 and 5.5, therefore we cannot exclude that differences might occur at these distinct pH values if residing for extensive period of time. However, previous studies in other biological models, showed that pH 5.5 or even 4.5 were not altering Mtb pH homeostasis and survival suggesting the Mtb can adapt and survive within these conditions (Darby et al., 2013; Fontes et al., 2020; Vandal et al., 2008). We also anticipate our system can be applied to other intracellular bacteria that temporarily inhabit acidic compartments such as *Salmonella* spp., *Shigella* spp. or *Coxiella burnetti*.

Our dual imaging allowed us to define if, in macrophages, the localisation in acidic compartments affected the intrabacterial pH after treatment with antibiotics used in the clinic to treat tuberculosis. Remarkably, out of four antibiotics, each with different modes of action, we found that in human macrophages, only PZA disrupts Mtb intrabacterial pH homeostasis in a concentration and time dependent manner. For BDQ, which is effective against Mtb within infected human macrophages (Giraud-Gatineau et al., 2020; Greenwood et al., 2019), we could confirm that treatment targets bacteria to acidic compartments (Giraud-Gatineau et al., 2020), however without resulting in significant changes in intrabacterial pH, as was previously suggested as the BDQ mode of action (Hards et al., 2018). This also suggests that intrabacterial pH imaging is not a reliable proxy of bacterial viability (at least in this system). Our findings seem to be different from what has been previously observed with *M. smegmatis* where it was proposed that global antibiotic-induced pH alterations should be considered a potential mechanism contributing to antibiotic efficacy (Bartek et al., 2016).

The study of the PZA molecular mechanism(s) of action, and its extensive association with acidic pH for efficacy, have been mostly performed *in vitro* (Mc and Tompsett, 1954; Zhang et al., 2002; Zhang et al., 1999; Zhang et al., 2013). Here, we provide compelling evidence that PZA acts as a bacterial protonophore that disrupts Mtb intrabacterial pH homeostasis *in cellulo*. These changes in intrabacterial pH homeostasis can be prevented by inhibiting the macrophage v-ATPase activity with ConA, likely impairing the conversion of POA^(-)^ into HPOA inside endolysosomes. The significant effect of ConA in preventing the pH-disruption correlates with an increase in bacterial growth showing that there is a link between pH-disruption and efficacy of protonophores as previously shown (Darby et al., 2013). This link between intrabacterial pH alteration and efficacy seems to apply only to PZA, since the other three very potent antibiotics we tested BDQ, RIF and INH are also effective against intracellular Mtb without affecting the intrabacterial pH.

Finally, by combining our dual imaging approach with mycobacterial mutants and naturally PZA-resistant strains, we were able to define if intracellular localisation affects the pH-disruption related efficacy of PZA. Notably two different mutants with a deficient ESX-1 secretion system showed a substantial reduction in growth when compared to the wild type strain after PZA treatment, a finding that reflects the increase in PZA accumulation reported previously (Santucci et al., 2021). Indeed, NanoSIMS analysis of PZA/POA accumulation showed that two-times more ΔRD1 bacteria were positive for this antibiotic in comparison to the WT strain (Santucci et al., 2021), which supports the two-fold difference in PZA EC_50_ we observed in this study. Similarly, ConA treatment negatively impacted by 3-4 times the amount of WT bacteria displaying a positive signal for the antibiotic by NanoSIMS (Santucci et al., 2021), which was reflected by the 3.5-fold difference in antibacterial efficacy when analysing the EC_50_ of PZA alone or in combination with ConA in the present study. Such results suggest a strong correlation between antibiotic accumulation and antibacterial efficacy as previously reported using bulk LC-MS/MS analysis (Richter et al., 2017). The use of mutants with different lifestyles in our experimental system has highlighted that cytosolic localisation can be an important factor that dictates antibiotic susceptibility *in cellulo*. Our data with ESX-1 defective mutants suggest that continuous and homogenous residence in a phagosome affects sensitivity to PZA and subsequently intrabacterial pH homeostasis disruption.

Within macrophages, Mbv was not sensitive to PZA which did not affect intrabacterial pH suggesting that POA, but not PZA, is primarily responsible for the pH-disruption effect. This confirms that POA is the main active form of the drug and that its inhibitory effect is tightly linked to pH *in cellulo*. Importantly, our results show that PZA accumulation and efficacy require acidic environments within host cells, highlighting that the pH-dependent mechanism of action is crucial in more sophisticated biological systems (Lamont et al., 2020). It is worth mentioning that the results obtained in this study are different from the one described *in vitro* by Peterson *et al*. (Peterson et al., 2015) showing that PZA/POA did not induce pH homeostasis disruption even at pH 5.8. These discrepancies could be explained by some differing experimental parameters including differences between *in cellulo* and *in vitro* investigations, the use of H37Ra and H37Rv strains, and the increased sensitivity of fluorescence microscopy in contrast to bulk spectrofluorimetric analysis. Finally, more investigations will be required to determine whether PanD inhibition could be potentially targeted by the drug under these conditions, and directly or indirectly involved in this pH-dependent antibacterial inhibition.

Because wild type Mtb displays a heterogenous subcellular localisation, where a fraction of Mtb is localised in phagosomes but another fraction is in the cytosol, we postulate that dynamic and heterogenous environments contribute to the pH-disruptive action of PZA in human macrophages. We believe that such methodology, our findings, and the concepts that have emerged from this study will be valuable to characterise antibiotic modes of action *in cellulo*.

### Significance

We still do not completely understand why tuberculosis treatment requires the combination of several antibiotics for up to six months. Mtb is an intracellular pathogen and it is still unknown whether heterogenous and dynamic intracellular populations of bacteria in different cellular environments affect antibiotic efficacy. By developing a dual live imaging approach to monitor mycobacterial pH homeostasis, host-cell environment and antibiotic action, we show here that intracellular localisation of Mtb affects the efficacy of one first-line anti-TB drug. Our observations can be applicable to the treatment of other intracellular pathogens and help to inform the development of more effective combined therapies for tuberculosis that target heterogenous bacterial populations within the host.

## Abbreviations

(TB): tuberculosis
(Mtb): *Mycobacterium tuberculosis*
(RIF): rifampicin
(INH): isoniazid
(EMB): ethambutol
(PZA): pyrazinamide
(POA): total pyrazinoic acid
(POA^(-)^): pyrazinoate anion
(HPOA): neutral protonated pyrazinoic acid
(CLEIM): correlative light, electron, ion microscopy
(MDM): human monocyte-derived macrophages
(iPSC): human induced pluripotent stem cells
(iPSDM): human induced pluripotent stem cell-derived macrophages
(ConA): Concanamycin A
(mROI): mycobacterial region of interest
(EC_50_): half maximal effective concentration
(BDQ): bedaquiline
(Mbv): *Mycobacterium bovis*
(MOI): multiplicity of infection

## Supplemental Information

Supplemental Information is attached to this manuscript and contains 6 additional figures with their corresponding legends.

## Acknowledgements

We would like to acknowledge all members of the Host-Pathogen Interactions in Tuberculosis laboratory for insightful discussions. We thank D. Branch Moody for critical reading of the manuscript and helpful discussions. This work was supported by the Francis Crick Institute (to MGG), which receives its core funding from Cancer Research UK (FC001092), the UK Medical Research Council (FC001092), and the Wellcome Trust (FC001092). This project has also received funding from the European Research Council (ERC) under the European Union’s Horizon 2020 research and innovation programme (grant agreement n°772022). For the purpose of Open Access, the authors have applied a CC BY public copyright licence to any Author Accepted Manuscript version arising from this submission. CB has received funding from the European Respiratory Society and the European Union’s H2020 research and innovation programme under the Marie Sklodowska-Curie grant agreement n°713406. PS is supported with a non-stipendiary FEBS long-term fellowship and has received funding from the European Union’s H2020 research and innovation programme under the Marie Sklodowska-Curie grant agreement *SpaTime_AnTB* n°892859.

## Authors contribution

PS and MGG conceived the study and designed the experiments. PS performed most of the experiments with help from BA, LB, EB, CB, EP and NA. All authors provided intellectual input by analysing and/or discussing data. PS and MGG wrote the manuscript. All authors read the manuscript and provided critical feedback before submission.

## STAR ★ Methods

### Key Resources Table

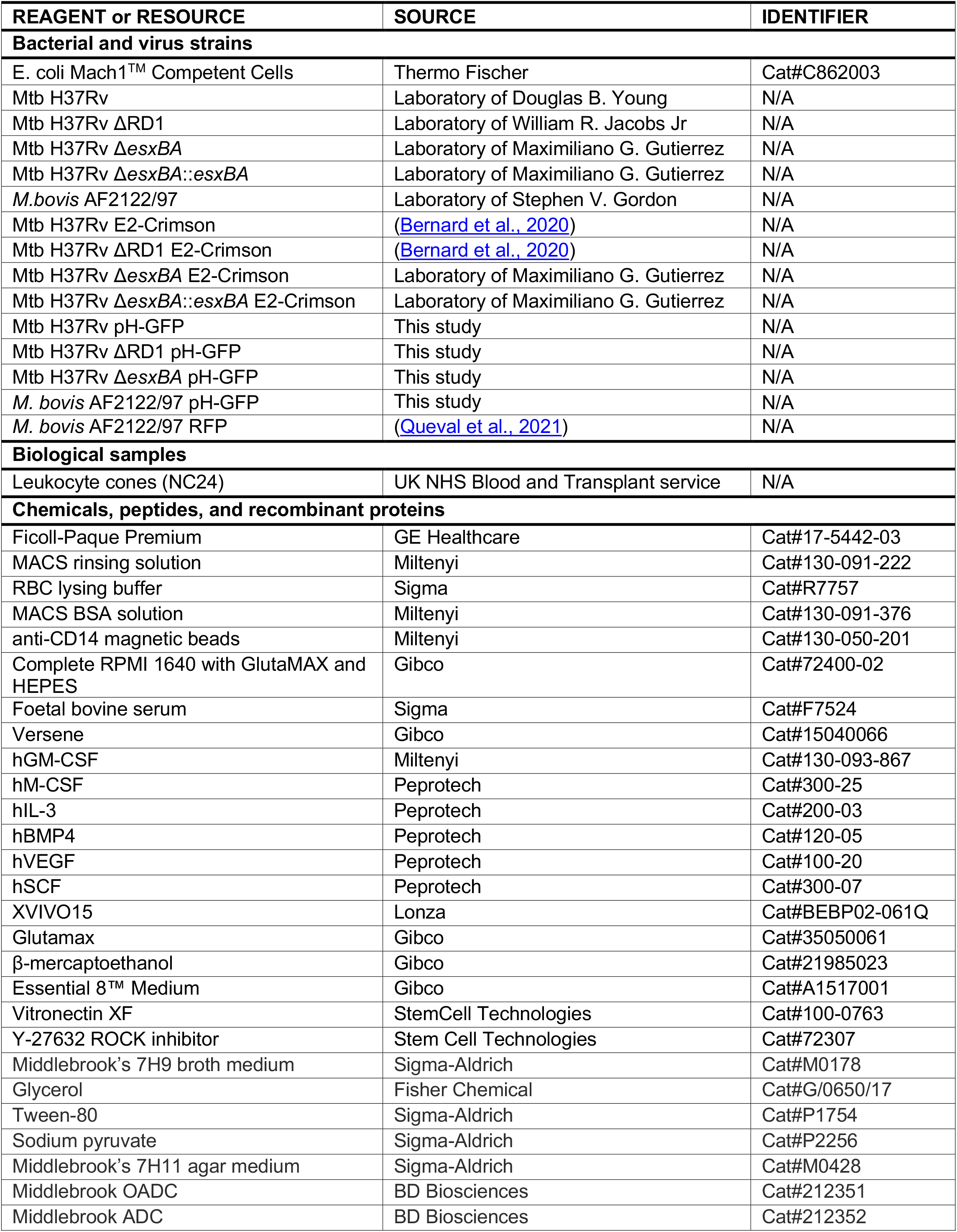
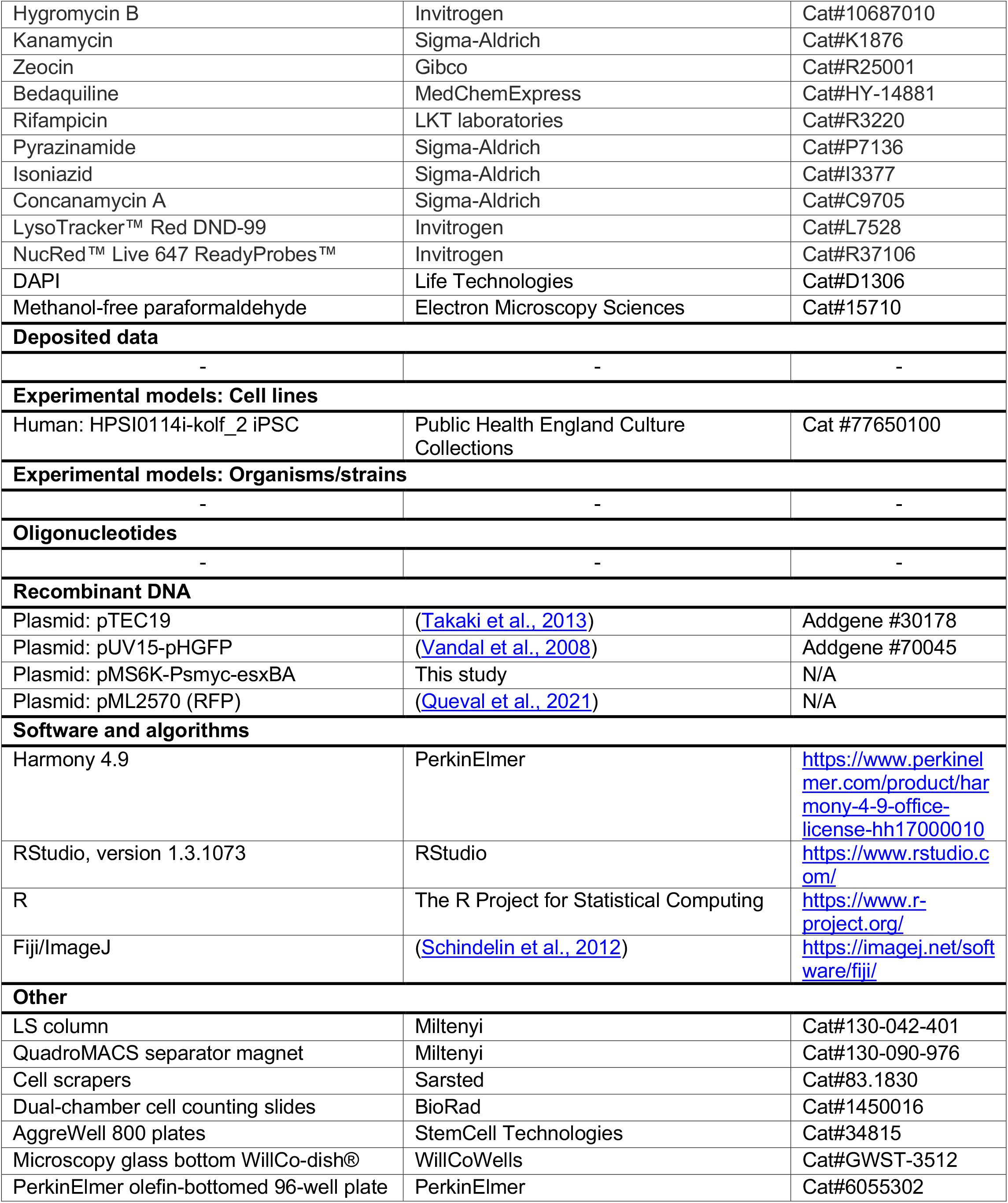

## Resource availability

### Lead contact

Further information and requests for resources and reagents should be directed to and will be fulfilled by the lead contact, Maximiliano G. Gutierrez.

### Materials availability

All unique or stable reagents generated in this study are available from the lead contact upon reasonable request. Availability of materials might be subjected to Materials Transfer Agreement (MTA) establishment.

### Data and code availability

All data reported in this paper will be shared by the lead contact upon reasonable request. This paper does not report original code.

## Experimental model and subject details

### Mycobacterial strains and culture conditions

*Mycobacterium tuberculosis* (Mtb) H37Rv and ΔRD1 strains were obtained from William R. Jacobs Jr. (Albert Einstein College of Medicine, New-York, USA), Suzie Hingley-Wilson (University of Surrey, Guilford, UK) and Douglas B. Young (The Francis Crick Institute, London, UK). *Mycobacterium bovis* AF2122/97 (Mbv) reference strain was provided Stephen V. Gordon (University College Dublin, Dublin, Ireland). Mtb Δ*esxBA* mutant was generated in our laboratory by using the ORBIT system, genetically mapped by PCR and sequenced (Aylan *et al*., *in preparation*). Its respective complement Mtb Δ*esxBA*::*esxBA* was generated by transformation with a mycobacterial kanamycin resistant integrative vector carrying a functional copy of *esxBA* genes under the control of the Psmyc promoter (Aylan *et al*., *in preparation*). Both clones did not show any growth impairment *in vitro* and were validated based on ESAT-6 and CFP-10 production and secretion by conventional immunoblot (Aylan *et al*., *in preparation*). Recombinant Mtb or Mbv strains expressing pH-GFP (pUV15-pHGFP; Addgene Plasmid #70045, kindly gifted by Sabine Ehrt), RFP (pML2570) or E2-Crimson (pTEC19, Addgene Plasmid #30178, kindly gifted by Lalita Ramakrishnan) fluorescent proteins were generated by electroporation and further selected onto appropriate medium. Recombinant Mtb strains were grown in Middlebrook 7H9 broth supplemented with 0.2% glycerol (v/v) (Fisher Chemical, G/0650/17), 0.05% Tween-80 (v/v) (Sigma-Aldrich, P1754) and 10% ADC (v/v) (BD Biosciences, 212352) whereas recombinant Mbv strains expressing pH-GFP or RFP fluorescent protein were grown in 7H9 Middlebrook supplemented with 40 mM sodium pyruvate (Sigma-Aldrich, P2256). Bacterial cultures (10 mL) were incubated under constant rotation in 50 mL conical tubes at 37°C. Hygromycin B (Invitrogen, 10687010), kanamycin (Sigma-Aldrich, K1876) or zeocin (Invivogen, ant-zn-05) were used as a selection marker for the fluorescent strains at a concentration of 50 mg/L, 25 mg/L and 25 mg/L respectively. All selected clones were tested for PDIM positivity by thin layer chromatography of lipid extracts from cultures prior to performing infection experiments.

### Preparation and culture of human-monocyte derived macrophages

Human monocyte-derived macrophages (MDM) were prepared from Leukocyte cones (NC24) supplied by the NHS Blood and Transplant service as previously described (Greenwood et al., 2019; Lerner et al., 2017; Santucci et al., 2021). Briefly, white blood cells were isolated by centrifugation on Ficoll-Paque Premium (GE Healthcare, 17-5442-03) for 60 min at 300 g. Mononuclear cells were collected and washed twice with MACS rinsing solution (Miltenyi, 130-091-222). Cells were subsequently incubated with 10 mL RBC lysing buffer (Sigma, R7757) at room temperature. After 10 min, cells were washed with rinsing buffer and then were re-suspended in 80 µL MACS rinsing solution supplemented with 1% BSA (Miltenyi, 130-091-376) (MACS/BSA) and 20 µL anti-CD14 magnetic beads (Miltenyi, 130-050-201) per approximately 10^8^ cells. After 20 min at 4°C, cells were washed in MACS/BSA solution and re-suspended at a concentration of 2.10^8^ cells/mL in MACS/BSA and further passed through a pre-equilibrated LS column (Miltenyi, 130-042-401) in the field of a QuadroMACS separator magnet (Miltenyi, 130-090-976). The LS column was washed three times with MACS/BSA solution, then CD14 positive cells were eluted, centrifuged and re-suspended in complete RPMI 1640 with GlutaMAX and HEPES (Gibco, 72400-02), 10% foetal bovine serum (Sigma, F7524) containing 10 ng/ml of hGM-CSF (Miltenyi, 130-093-867). Differentiation was performed by plating approximately 10^6^ cells/mL in untreated petri dishes and further incubated in a humidified 37°C incubator with 5% CO_2_. After three days, an equal volume of fresh complete media including hGM-CSF was added. Six days after the initial isolation, differentiated macrophages were detached in 0.5 mM EDTA in ice-cold PBS using cell scrapers (Sarsted, 83.1830), pelleted by centrifugation and re-suspended in complete RPMI 1640 medium containing 10% foetal bovine serum where cell count and viability was estimated (BioRad, TC20™ Automated Cell Counter) before plating for experiments.

### Human-induced pluripotent stem cells culture and human induced pluripotent stem cells-derived macrophages preparation

Human iPSC maintenance and IPSDM preparation was performed as recently reported (Bernard et al., 2020). Briefly, KOLF2 IPSC (HPSI0114i-kolf_2 iPSC, Public Health England Culture Collections, Cat #77650100) were maintained in Vitronectin XF (StemCell Technologies, #100-0763) coated plates with E8 medium (ThermoFisher Scientific, A1517001) in a humidified 37°C incubator with 5% CO_2_. Cells were passaged by performing a 1/6 dilution when reaching approximately 70% confluency using Versene (Gibco, 15040066). Monocyte factories were set up following a previously reported protocol (van Wilgenburg et al., 2013). A single cell suspension of iPSC was generated in E8 medium containing 10 µM Y-27632 ROCK inhibitor (Stem Cell Technologies, # 72307) and seeded into AggreWell 800 plates (StemCell Technologies, # 34815) with approximately 4×10^6^ cells/well and centrifuged at 100 g for 5 min. The forming embryonic bodies (EB) were fed daily with two 50 % medium changes with E8 medium supplemented with 50 ng/ml hBMP4 (Peprotech, 120-05ET), 50 ng/ml hVEGF (Peprotech, 100-20) and 20 ng/ml hSCF (Peprotech, 300-07) for 3 days. On day 4, the EBs were harvested by flushing out of the well with gentle pipetting and filtered through an inverted 40 µm cell strainer. EBs were seeded at 250–300 per T225 flask in factory medium consisting of X-VIVO 15 (Lonza, BE02-061Q) supplemented with Glutamax (Gibco, 35050061), 50 µM β-mercaptoethanol (Gibco, 21985023), 100 ng/ml hM-CSF (Peprotech, 300-25) and 25 ng/ml hIL-3 (Peprotech, 200-03). Monocyte factories were fed once per week with factory medium for 4-5 weeks until monocyte production was observed in the supernatant. Up to 50% of the supernatant was harvested weekly and factories fed with 20-30 ml factory medium. For differentiation, the supernatant was centrifuged and cells resuspended in X-VIVO 15 supplemented with Glutamax and 20 ng/ml hGM-CSF and plated at 12×10^6^ cells per 15 cm petri dish to differentiate over 7 days, where a 50% medium change was performed on day 4. Seven days after the initial plating, differentiated macrophages were detached with Versene (Gibco, 15040066) for 15 min at 37°C and 5% CO_2_. Versene was further diluted 1:3 with PBS and cells were gently detached with cell scrapers (Sarsted, 83.1830), pelleted by centrifugation and re-suspended in X-VIVO 15 plus Glutamax where cell count and viability were estimated (BioRad, TC20™ Automated Cell Counter) before plating for experiments.

## Methods details

### Macrophage infection with Mtb and Mbv strains

For macrophage infection, mycobacterial inoculum was prepared following a well-established procedure (Lerner et al., 2017; Schnettger et al., 2017). First, bacterial cultures were pelleted by centrifuging approximately 10 mL of mid-exponential phase cultures (OD_600nm_ = 0.6 ± 0.2) at 4000 rpm for 5 min. Pellets were washed twice in sterile PBS buffer (pH 7.4), then, an equivalent volume of sterile 2.5-3.5 mm autoclaved glass beads was added to individual pellet and bacterial clumps were disrupted by vigorously shaking. Bacteria were re-suspended in the appropriate cell culture media and the clumps were removed by slow-speed centrifugation at 1200 rpm for 5 min. The supernatant containing the bacterial suspension of interest was transferred to a fresh tube and OD_600nm_ was measured to determine bacterial concentration. In this protocol, it was assumed that an OD_600nm_ of 1 approximates to 10^8^ bacteria/mL. For high-content dual imaging experiments and intracellular antibiotic assays, macrophages were infected with mycobacterial strains at a multiplicity of infection (MOI) of 1 for 2 h at 37°C. After 2 h of uptake, cells were washed with PBS to remove extracellular bacteria and fresh media was added.

### High-content dual-live fluorescence imaging, determination of Mtb intrabacterial pH-GFP fluorescence ratio and Mtb-associated LysoTracker intensity

For high-content live-cell imaging, cells were infected with fluorescent mycobacteria producing ratiometric pH-GFP at a MOI of 1 as described above. After, 24 hours the culture media was replaced by fresh medium only or fresh medium containing 50 nM ConA (Sigma-Aldrich, C9705) for IPSDM and 100 nM ConA for MDM. After 24 hours, infected cells were washed once with PBS buffer (pH 7.4) and stained with complete medium containing 200 nM LysoTracker™ Red DND-99 (Invitrogen, L7528) in a humidified 37°C incubator with 5% CO2. After 30 min, staining medium was removed and replaced with fresh medium containing NucRed™ Live 647 ReadyProbes™ (Invitrogen, R37106) following the manufacturer recommendations to facilitate cell detection. Live-cell imaging was further performed using the OPERA Phenix microscope with a 63x water-immersion objective. Image acquisition was performed with the confocal mode using the default autofocus function and a binning of 1. Mtb pH-GFP signal was detected using λex 405 nm/λem 500-550 nm and λex 488nm/λem 500-550 nm, LysoTracker signal was detected λex 561 nm/λem 570-630 nm and NucRed Live signal λex 640 nm/λem 650-760 nm. Laser power for all channels were set between 20% and 30% with an exposure time of 200 ms. Each channel was imaged independently and a minimum of 3 to 4 distinct focal z-planes spaced with 0.5-1 µm were acquired. Multiple fields of view (323 µm × 323 µm) from each individual well were imaged with a set overlap of 10 % in between fields. Segmentation and analysis were performed using the Harmony software (Perkin Elmer, version 4.9). Briefly, cellular region was detected based on the fluorescent signals in the far-red emission channel using the “Find Image Region” building block and the “Absolute Threshold” function. Intracellular Mtb pH-GFP were detected based on the GFP signal obtained into both λex 405 nm/λem 500-550 nm and λex 488nm/λem 500-550 nm channels using the “Find Image Region” building block and the “Absolute Threshold” function. Signal from both GFP channels were merged using the “Calculate Image” building block and the function “By Formula” were a channel A + B operation was applied. This combined image was filtered to reduce background noise using the “Filter Image” building block and a sliding parabola function. This Mtb mask was used to quantify Mtb pH-GFP mean fluorescent signal per single object from both 405nm/510nm and 488nm/510nm channels. Ratiometric signal were obtained by dividing the mean intensity quantified at λex 405 nm/λem 500-550 nm by the one obtained at λex 488nm/λem 500-550 nm for each object. To quantify Mtb-associated LysoTracker intensity, the Mtb mask was slightly extended using the “Find Surrounding Region” building block using the Method A with an individual threshold value of 0.8 and conservation of the input region. When assessing the spatiotemporal mode of action of PZA, human macrophages were infected for 24 hours and further treated with increasing concentration of PZA ranging from 0 to 400 mg/L in the absence or presence of ConA for additional 4-, 16-, 24-or 72 hours before acquisition was performed as describe above. When assessing anti-TB drug-mediated pH-disruption, Mtb-infected cells were left untreated or pulsed with PZA (100 mg/L), BDQ (2.5 mg/L), INH (5 mg/L) or RIF (5 mg/L) for 24 hours before imaging. Determination of absolute changes of pH-GFP ratio relative to the control condition (also referred as Δintrabacterial pH) was done by subtracting the value obtained in each experimental condition to its corresponding control condition. All the results were exported as CSV files, imported in the R studio software (The R Project for Statistical Computing, version 1.3.1073) and most of the graphs, displayed as boxplot, scatter plots or raincloud plots, were plotted with the ggplot2 package (version 3.3.2).

### Low-content dual-live fluorescence imaging and live-cell imaging

For low-content dual imaging and live-cell imaging, experimental set up and acquisition was performed as previously described (Bernard et al., 2020; Schnettger and Gutierrez, 2017; Schnettger et al., 2017) with slight modifications. Approximately 1 × 10^6^ IPSDM were seeded within 12 mm aperture glass bottom dishes (WillCo-dish®, GWST-3512) in 1 mL of X-VIVO 15 media plus Glutamax. Adherent cells were infected with fluorescent Mtb producing ratiometric pH-GFP at a MOI of 1 as described above and after 24 hours of infection medium was replaced with complete medium containing 200 nM LysoTracker™ Red DND-99 (Invitrogen, L7528). The dish was placed on a custom-made 35 mm dish holder and further incubated in a humidified 37°C incubator with 5% CO_2_. After 30 min of staining, dish was set under a Leica SP5 laser scanning confocal microscope (Leica Biosystems) in an environmental control chamber providing 37°C, 5 % CO_2_ and 20–30 % humidity for an additional 1 hour to avoid drifting issues upon acquisition. Image acquisition was performed with a HC PL APO CS2 63×/1.40 oil objective. Images of 1024 × 1024 pixels were acquired with Diode 405 nm, Argon 488 nm and DPSS 561 nm lasers where intensities were set up as 2%, 8% and 8% respectively. Emitted signal was collected at λem 510 ± 30 nm and λem 585 ± 15 nm for pH-GFP and LysoTracker channels respectively. One single Z-plane was acquired for each field and a minimum of 5 fields per biological sample were imaged. Determination of the Mtb pH-GFP ratio and Mtb-associated LysoTracker mean intensity values were performed by manual quantification as previously described (Bernard et al., 2020; Santucci et al., 2021). Briefly, the mROI were duplicated, the bacteria containing channel was manually thresholded and a single ‘Dilate’ command was applied to generate a binary mask in Fiji corresponding to the bacteria surrounded by one single ring of pixel. This mask was then used to measure the mean fluorescence intensity of pixels in λex 405 nm/λem 510 ± 30 nm, λex 488nm/λem 510 ± 30 nm and λex 510 nm/λem 585 ± 15 nm channels using the command ‘Measure’.

For live-cell imaging a very similar experimental set up was used where images were acquired every 30 min intervals over a time frame of 6 h to minimise photobleaching and phototoxicity. Same settings were used and 16 z-stacks of approximately 500nm were performed to catch most of the events contained within Mtb-infected cells. mROI were defined and tracked manually by selecting the appropriate focal plane over the course of the kinetics as previously described. Selected planes were then combined together using the ‘Concatenate’ command in Fiji (Schnettger and Gutierrez, 2017). Determination of Mtb pH-GFP and its respective LysoTracker-associated mean intensity was performed as mentioned above and further analysed overtime. All the results were exported as CSV files, imported in the R studio software (The R Project for Statistical Computing, version 1.3.1073) and the graphs were plotted with the ggplot2 package (version 3.3.2).

### Intracellular replication assays in Mtb- and Mbv-infected macrophages

Intracellular replication assays were performed by high-content fluorescence quantitative imaging as previously described (Greenwood et al., 2019; Santucci et al., 2021). Briefly, 3.5-4.0 × 10^4^ cells per well were seeded into an olefin-bottomed 96-well plate (Perkin Elmer, 6055302) 16-20 hours prior to infection. Cells were infected as described above with pH-GFP or E2-Crimson producing strains for 24 hours and the culture media was replaced by fresh media containing increasing concentrations of PZA, RIF, INH, BDQ or left untreated. When indicated, fresh medium containing 50 nM ConA for IPSDM and 100 nM ConA for MDM (Sigma-Aldrich, C9705) was added together with the antibiotics. At the required time points, infected cells were washed with PBS buffer (pH 7.4) and fixed with a 4% methanol-free paraformaldehyde (Electron Microscopy Sciences, 15710) in PBS buffer (pH 7.4) for 16-20 h at 4°C. Fixative was removed and cells were washed in PBS buffer (pH 7.4) before performing the appropriate nuclear staining using either DAPI (Invitrogen, D1306) or NucRed™ Live 647 ReadyProbes™ (Invitrogen, R37106) for nuclear visualisation. Image acquisition was performed with the OPERA Phenix high-content microscope with a 40x water-immersion 1.1 NA objective. The confocal mode with default autofocus and a binning of 1 was used to image multiple fields of view (323 µm × 323 µm) from each individual well with 10% overlapping, where acquisition was performed at 4 distinct focal planes spaced with 1 or 2 µm. Imaging of stained nuclei and fluorescent bacteria was done with similar λex/λem settings as described above. Analysis was performed using the Harmony software (Perkin Elmer, version 4.9) where maximum projection of the 3-4 z-planes was used to perform single cell segmentation by using the “Find nuclei” and “Find cells” building blocks. Cells on the edges were excluded from the analysis. The fluorescent bacterial signal was detected using the “Find Image Region” building block where a manual threshold was applied to accurately perform bacterial segmentation. The Mtb area per cell was determined by quantifying the total area (expressed in µm^2^) of GFP^+^ or E2-Crimson^+^ signal per single macrophage. The relative growth index was determined by using the following formula ∼ (Mean Mtb area per cell t_96h_ - Mean Mtb area per cell t_24h_) / (Mean Mtb area per cell t_24h_) and the relative values were obtained by using the untreated control as a reference of 100% growth (0% inhibition). All the results were exported as CSV files, imported in the R studio software (The R Project for Statistical Computing, version 1.3.1073) and graphs were plotted with the ggplot2 package (version 3.3.2).

### Quantification and Statistical analysis

Results displayed were obtained from n = 2, n = 3 or n = 4 biologically independent experiments performed at least each time in two-three technical replicates (unless otherwise stated). The statistical tests used, the number of biologically independent replicates, the number of technical replicates and the number of single-cell or single-mROI analysed are indicated in each figure legend. Statistical analysis by pairwise comparison was performed using Wilcoxon signed-rank test with the ‘*wilcox*.*test()*’ function in R where differences were considered statistically significant when *p* ≤ 0.05. Statistical analysis is displayed in the figure as ^*^ *p* ≤ 0.05; ^**^ *p* ≤ 0.01; ^***^ *p* ≤ 0.001 or alternatively as ^#^ *p* ≤ 0.05; ^##^ *p* ≤ 0.01; ^*###*^ *p* ≤ 0.001. All the *p-*values contained in the text or the figures are relative to the control condition (unless otherwise stated). Spearman rank correlation coefficient (*r*_*s*_) and its corresponding *p*-value were calculated by using the ggpubr R package and assessed by two-tailed statistical *t*-test.

