## Supplementary information for "Visualizing pyrazinamide action by live single cell imaging of phagosome acidification and *Mycobacterium tuberculosis* pH homeostasis"

### Figure S1

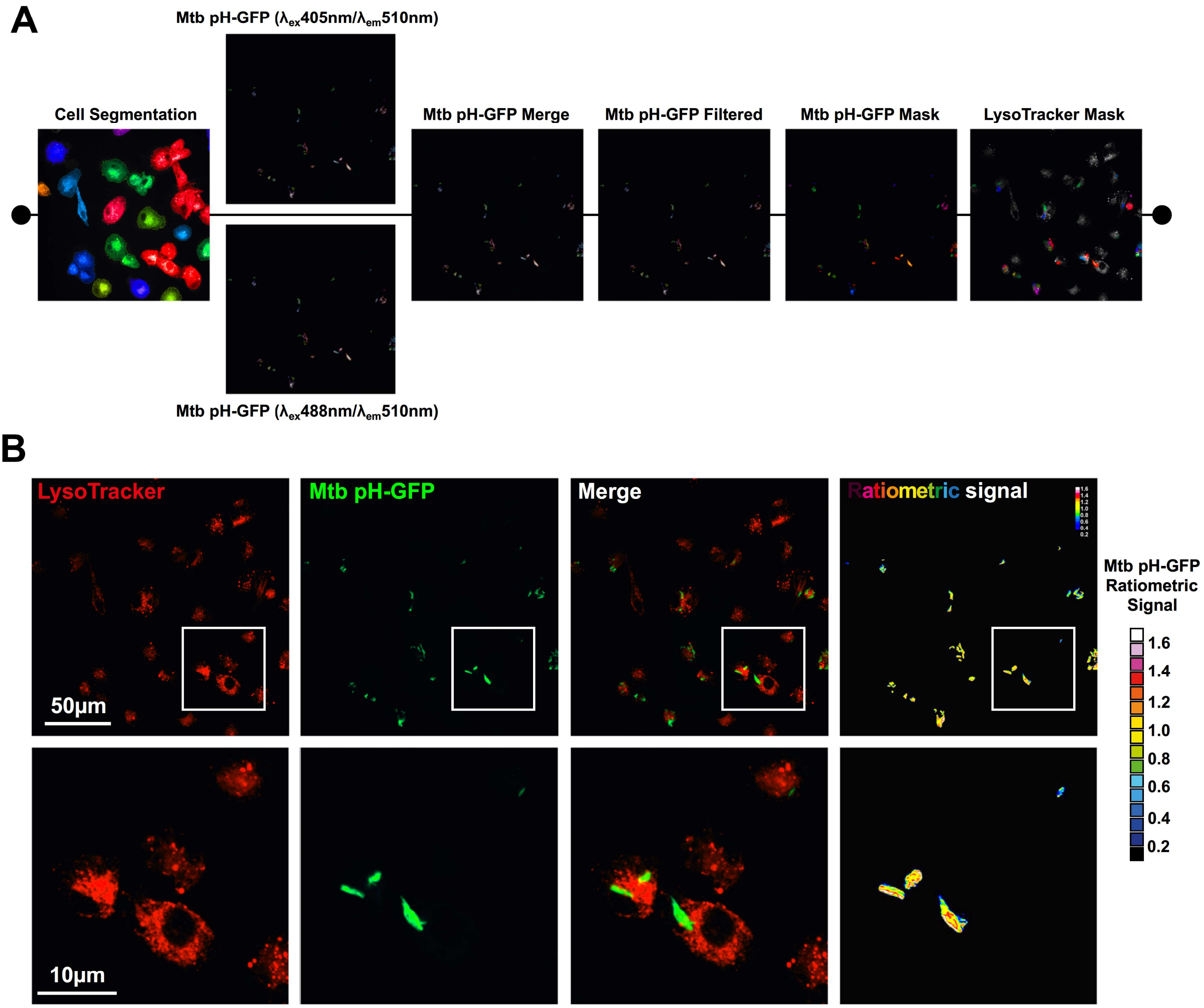
A
Mtb pH-GFP (λex405nm/λem510nm)
Cell Segmentation
Mtb pH-GFP Merge
Mtb pH-GFP Filtered
Mtb pH-GFP Mask
LysoTracker Mask
Mtb pH-GFP (λex488nm/λem510nm)
B
LysoTracker
Mtb pH-GFP
Merge
Ratiometric signal
50μm
10μm
Mtb pH-GFP Ratiometric Signal
1.6
1.4
1.2
1.0
0.8
0.6
0.4
0.2

### Figure S2

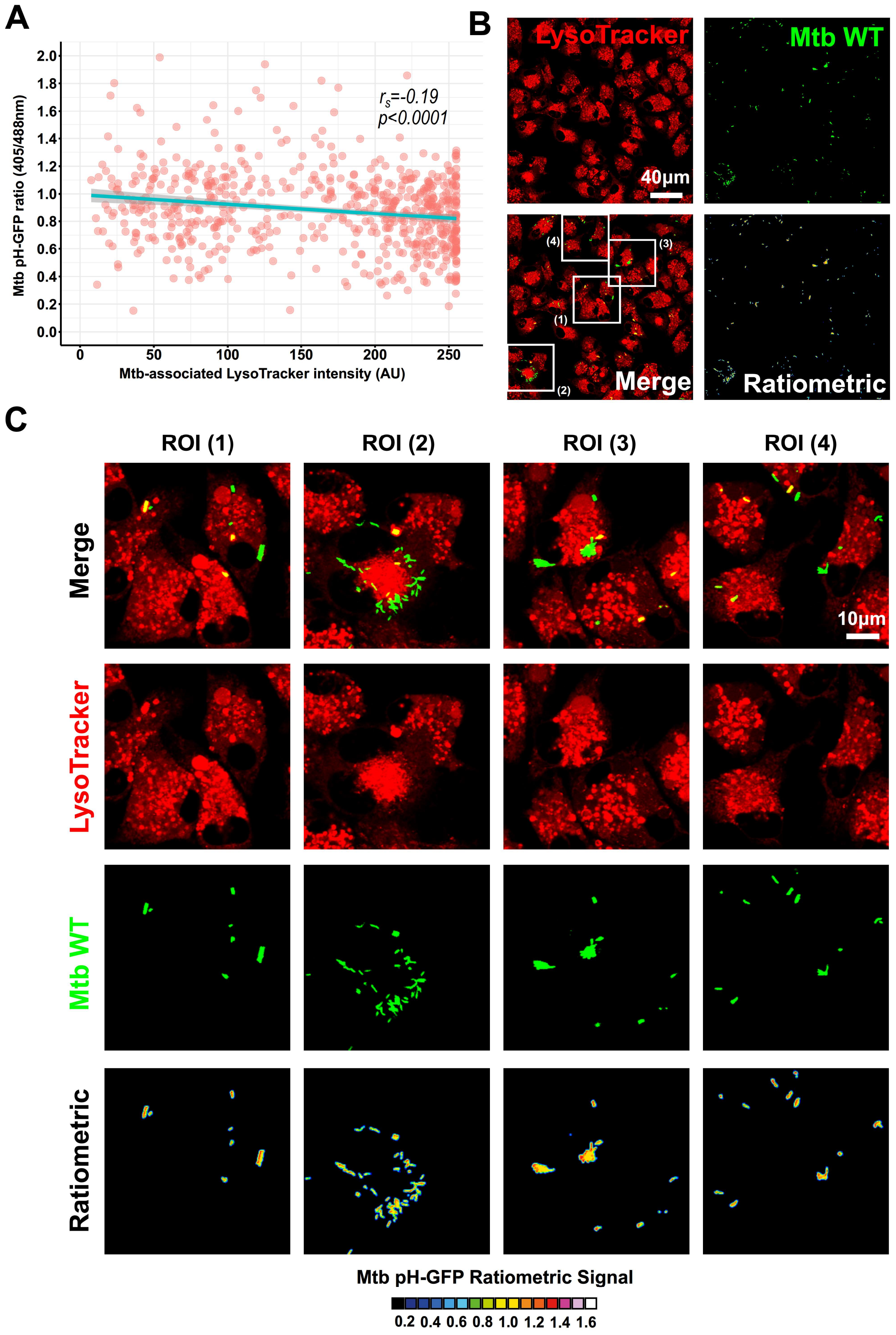
A
Mtb pH-GFP ratio (405/488nm)
2.0
1.8
1.6
1.4
1.2
1.0
0.8
0.6
0.4
0.2
0.0
0
50
100
150
200
250
Mtb-associated LysoTracker intensity (AU)
$r_s$=-0.19
$p$<0.0001
B
LysoTracker
Mtb WT
40μm
(1)
(2)
(3)
(4)
Merge
Ratiometric
C
ROI (1)
ROI (2)
ROI (3)
ROI (4)
Merge
LysoTracker
Mtb WT
Ratiometric
10μm
Mtb pH-GFP Ratiometric Signal
0.2 0.4 0.6 0.8 1.0 1.2 1.4 1.6

### Figure S3

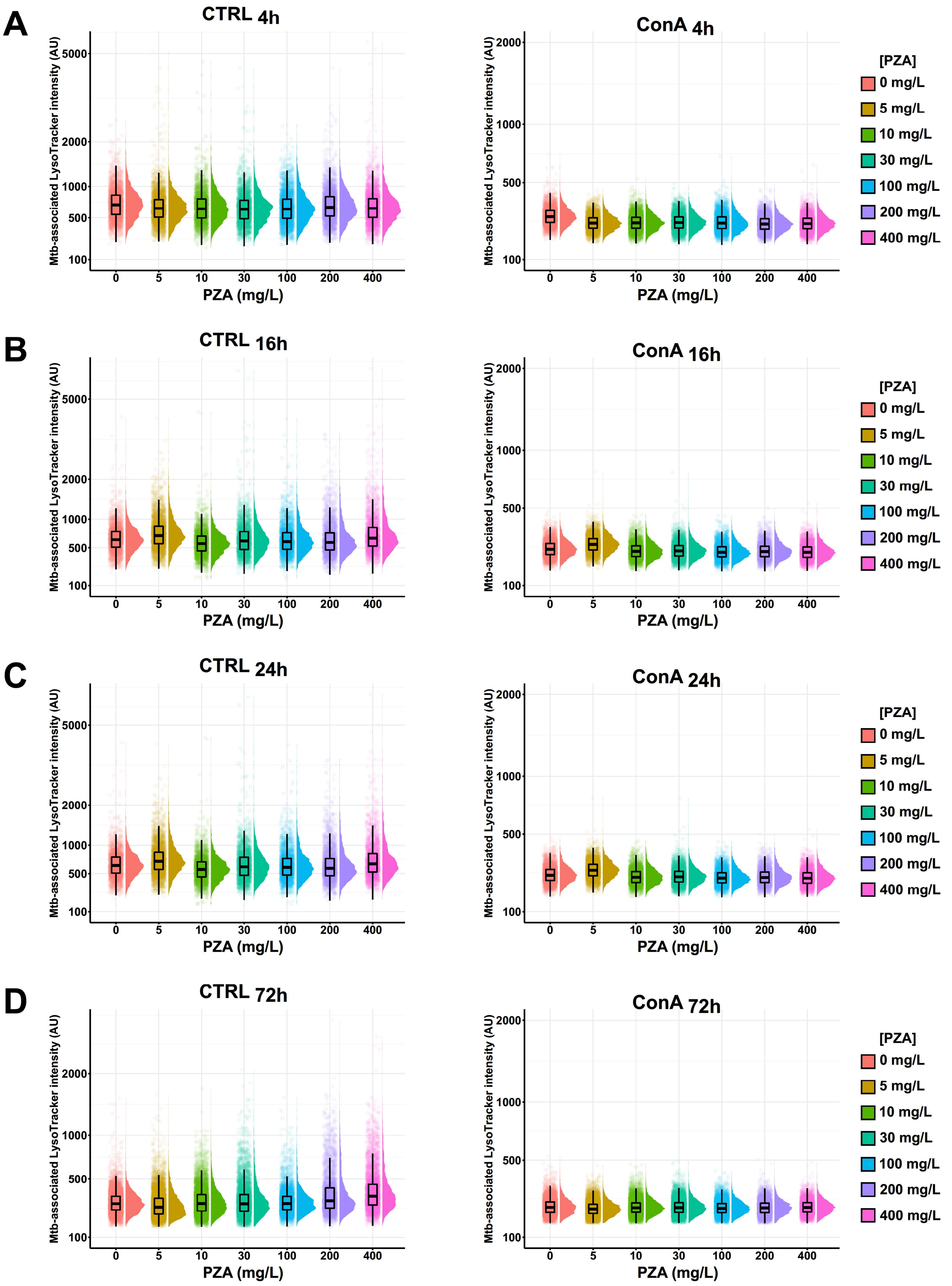

A
CTRL 4h
ConA 4h
B
CTRL 16h
ConA 16h
C
CTRL 24h
ConA 24h
D
CTRL 72h
ConA 72h
Mtb-associated LysoTracker intensity (AU)
PZA (mg/L)
0
5
10
30
100
200
400
[PZA]
0 mg/L
5 mg/L
10 mg/L
30 mg/L
100 mg/L
200 mg/L
400 mg/L

### Figure S4

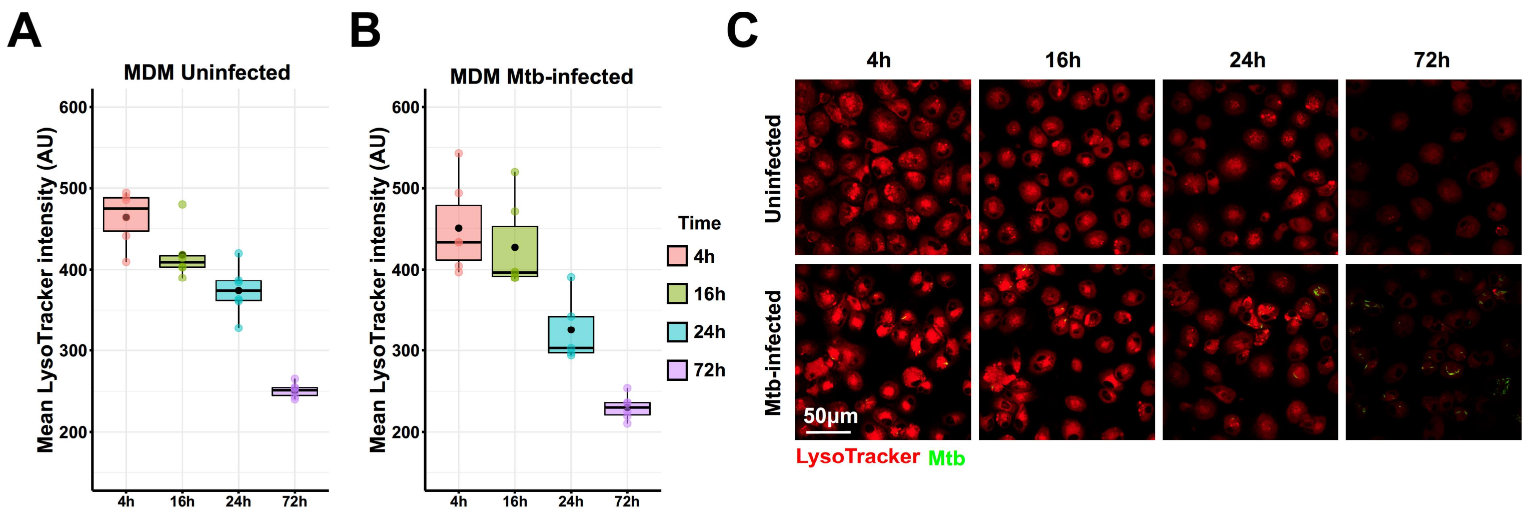

A
B
C
MDM Uninfected
MDM Mtb-infected
Mean LysoTracker intensity (AU)
600
500
400
300
200
4h
16h
24h
72h
Time
Uninfected
Mtb-infected
50μm
LysoTracker
Mtb

### Figure S5

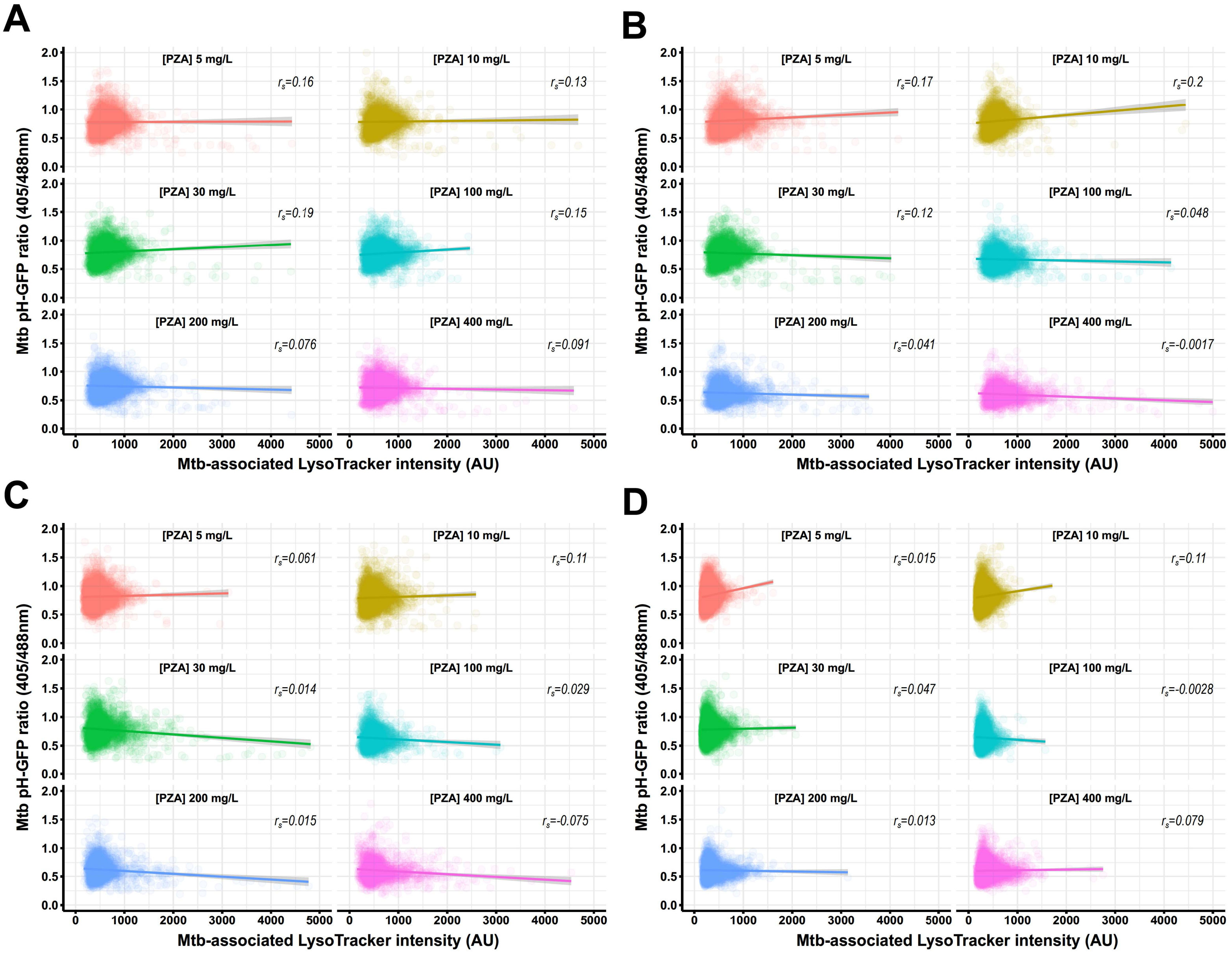
A
B
C
D
[PZA] 5 mg/L
[PZA] 10 mg/L
[PZA] 30 mg/L
[PZA] 100 mg/L
[PZA] 200 mg/L
[PZA] 400 mg/L
Mtb pH-GFP ratio (405/488nm)
Mtb-associated LysoTracker intensity (AU)
A: $r_s$=0.16, $r_s$=0.13, $r_s$=0.19, $r_s$=0.15, $r_s$=0.076, $r_s$=0.091
B: $r_s$=0.17, $r_s$=0.2, $r_s$=0.12, $r_s$=0.048, $r_s$=0.041, $r_s$=-0.0017
C: $r_s$=0.061, $r_s$=0.11, $r_s$=0.014, $r_s$=0.029, $r_s$=0.015, $r_s$=-0.075
D: $r_s$=0.015, $r_s$=0.11, $r_s$=0.047, $r_s$=-0.0028, $r_s$=0.013, $r_s$=0.079

### Figure S6

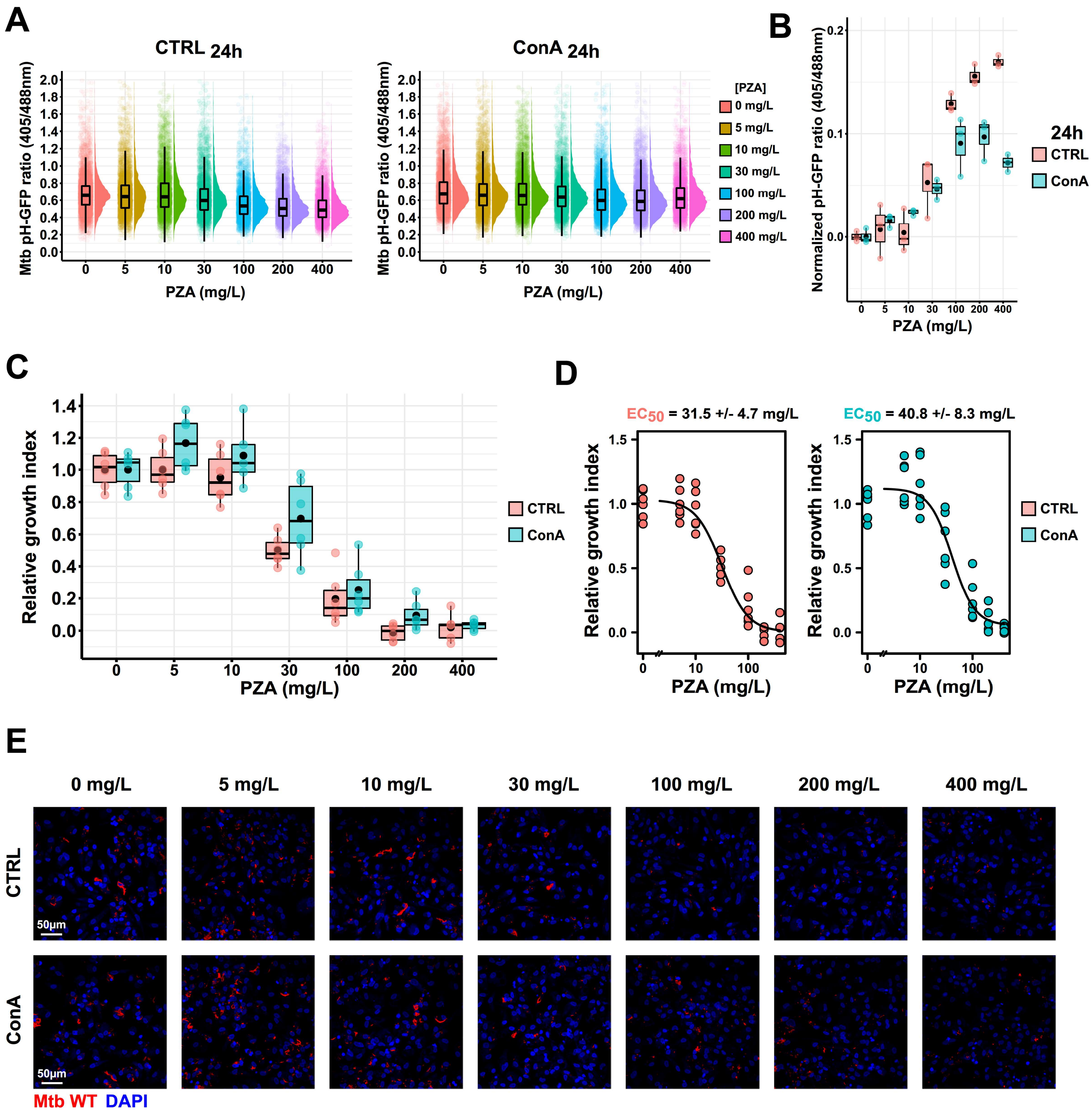
A
CTRL 24h
ConA 24h
Mtb pH-GFP ratio (405/488nm)
PZA (mg/L)
[PZA]
0 mg/L
5 mg/L
10 mg/L
30 mg/L
100 mg/L
200 mg/L
400 mg/L
B
Normalized pH-GFP ratio (405/488nm)
24h
CTRL
ConA
C
Relative growth index
D
EC50 = 31.5 +/- 4.7 mg/L
EC50 = 40.8 +/- 8.3 mg/L
E
0 mg/L
5 mg/L
10 mg/L
30 mg/L
100 mg/L
200 mg/L
400 mg/L
CTRL
ConA
50µm
Mtb WT DAPI

### *Supplemental information*

### Visualizing pyrazinamide action by live single cell imaging of phagosome acidification and *Mycobacterium tuberculosis* pH homeostasis

Pierre Santucci[1*], Beren Aylan[1], Laure Botella[1], Elliott M. Bernard[1#], Claudio Bussi[1], Enrica Pellegrino[1], Natalia Athanasiadi[1] and Maximiliano G. Gutierrez[1*§]

**Affiliations:**

[1] Host-Pathogen Interactions in Tuberculosis Laboratory, The Francis Crick Institute, 1 Midland Road, London, NW1 1AT, United Kingdom.

[#] Present Address: Department of Biochemistry, University of Lausanne, 1066 Epalinges, Switzerland

[§] Lead contact

**Figure S1: Analytical pipeline used in this study to perform high-content quantitative analysis of fluorescence microscopy images**

**(A)** Schematic representation of the segmentation and analysis pipeline used in this study to perform quantitative analysis of Mtb intrabacterial pH and Mtb-associated LysoTracker intensity by fluorescence microscopy. **(B)** Representative images of Mtb pH-GFP-infected MDMs stained with LysoTracker. Infected cells were infected for 48 hours then pulsed with 200 nM of LysoTracker Red for 30 min. Cells were imaged live using the high-content screening platform OPERA Phenix. Micrographs display LysoTracker labelling (red) and Mtb pH-GFP (green). Ratiometric signal was obtained by dividing the fluorescence intensity acquired with excitation/emission channels of 405/510 nm by the one obtained at 488/510 nm. Ratiometric signal is displayed as a 16-colours palette ranging from 0 to 1.6 units. Scale bar corresponds to 50 µm. Region of interests highlighted by the white rectangles, are shown in detail in the bottom panels respectively. Scale bar corresponds to 10 µm. The complete procedures are detailed in the *STAR ✦ Methods* section of this manuscript.

**Figure S2: Monitoring of Mtb intrabacterial pH and host-cell intracellular acidification by an alternative low-content imaging approach**

Human macrophages were infected for 48 hours and then pulsed with 200 nM of LysoTracker Red for 30 min before live-acquisition was performed using a Leica SP5 AOBS Laser Scanning Confocal Microscope. Quantitative analysis of Mtb pH-GFP ratio (405/488nm) and Mtb-associated LysoTracker were performed using the open source Fiji software. **(A)** Spearman's correlation between Mtb-associated LysoTracker (x-axis) and Mtb pH-GFP ratio (405/488nm) (y-axis) signals in individual bacterial regions of interest within infected-IPSDM. The linear regression is shown as the cyan line, the Spearman rank correlation coefficient ($r_s$) and the corresponding *p*-value were calculated by using the ggpubr R package and two-tailed statistical t-test. **(B)** Representative micrographs display LysoTracker labelling (red) and Mtb pH-GFP (green). Ratiometric signal was obtained by dividing the fluorescence intensity acquired with excitation/emission channels of 405/510 nm by the one obtained at 488/510 nm. Ratiometric signal is displayed as a 16-colours palette ranging from 0 to 1.6 units. Scale bar corresponds to 50 µm. **(C)** Regions of interest 1 to 4 from highlighted by the white rectangles in **(B)**, are shown in detail in the bottom panels respectively. Scale bar corresponds to 10 µm. Results are representative are from n = 2 biologically independent experiments performed at least in two-three technical replicates.

**Figure S3: Temporal dynamics of Mtb-associated LysoTracker intensity upon PZA-treatment *in cellulo***

Human macrophages were infected for 24 hours and subsequently treated with increasing concentration of PZA ranging from 0-400 mg/L in absence or presence of ConA for 4 h, 16 h, 24 h or 72 hours. Cells were then pulsed with 200 nM of LysoTracker Red for 30 min before live-acquisition was performed using the OPERA Phenix imaging platform. **(A-D)** Quantification of Mtb-associated Lysotracker mean intensity within infected-MDM treated with increasing concentration of PZA ranging from 0-400 mg/L in the absence or presence of v-ATPase inhibitor ConA at 4 h, 16 h, 24 h or 72 hours post-treatment. Results are displayed as raincloud plots where, black box-plots are overlaid on top of individual raw data and associated with their respective density plots. Each colour represents a specific PZA concentration. Results are from n = 2 biologically independent experiments performed at least in two-three technical replicates.

**Figure S4: Temporal dynamics of LysoTracker intensity within infected and uninfected human macrophages**

Human macrophages were infected with Mtb pH-GFP for 24 hours or left uninfected for an additional 4 h, 16 h, 24 h or 72 hours. Cells were then pulsed with 200 nM of LysoTracker Red for 30 min before live-acquisition was performed using the OPERA Phenix imaging platform. **(A-B)** Quantification of LysoTracker mean intensity within **(A)** uninfected MDM or **(B)** Mtb-infected MDM cellular region was determined at 4 h, 16 h, 24 h or 72 hours post-treatment. Results are displayed as box-plots with individuals' data. Black dots were added to highlight the mean of each conditions. Each colour represents a specific timepoint. **(C)** Representative micrographs display LysoTracker labelling (red) and Mtb pH-GFP (green). Scale bar corresponds to 50 µm. Results are from n = 2 biologically independent experiments performed at least in two-three technical replicates.

**Figure S5: PZA-mediated disruption of Mtb intrabacterial pH does not directly correlate with Mtb-asscociated LysoTracker intensity within human macrophages**

Human macrophages were infected for 24 hours and subsequently treated with increasing concentration of PZA ranging from 0-400 mg/L for 4 h, 16 h, 24 h or 72 hours. Cells were then pulsed with 200 nM of LysoTracker Red for 30 min before live-acquisition was performed using the OPERA Phenix imaging platform. **(A-D)** Spearman's correlation between Mtb-associated LysoTracker (x-axis) and Mtb pH-GFP ratio (405/488nm) (y-axis) signals in individual bacterial region of interests within infected-MDM at 4 h, 16 h, 24 h or 72 hours post-treatment respectively. Spearman rank correlation coefficient ($r_s$) shown as regression line was calculated by using the ggpubr R package. Each colour represents a specific PZA concentration. Results are from n = 2 biologically independent experiments performed at least in two-three technical replicates.

**Figure S6: PZA triggers Mtb intrabacterial pH-disruption in GM-CSF derived IPSDM**

Human macrophages were infected for 24 hours and subsequently treated with increasing concentration of PZA ranging from 0-400 mg/L in absence or presence of ConA for 24 hours. Cells were then pulsed with 200 nM of LysoTracker Red for 30 min before live-acquisition was performed using the OPERA Phenix imaging platform. **(A)** Quantification of Mtb pH-GFP ratio (405/488nm) within infected-IPSDM treated with increasing concentration of PZA ranging from 0-400 mg/L in the absence or presence of v-ATPase inhibitor ConA at 24 hours post-treatment. Results are displayed as raincloud plots where black box-plots are overlaid on top of individual raw data and associated with their respective density plots. Each colour represents a specific PZA concentration. **(B)** Determination of absolute changes in Mtb pH-GFP ratio (405/488nm) upon PZA treatment. Mean pH-GFP ratio of each conditions were subtracted to the PZA untreated condition in the absence or presence of ConA at 24 hours post-treatment to obtained an absolute value reflecting PZA-mediated pH disruption normalized to PZA free conditions. Determination of normalized pH-GFP ratio was performed in the absence (pink) or presence (cyan) of v-ATPase inhibitor ConA. Results are displayed as box-plots with individuals' raw data. Black dots were added to highlight the mean of each conditions. Results are representative from n = 2 biologically independent experiments performed at least in two-three technical replicates. **(C)** Quantitative analysis of E2-Crimson Mtb WT replication at the single cell level within IPSDM treated with increasing concentration of PZA in the absence or presence of ConA. Normalization was done to the mean Mtb area per cell pre-treatment ($t_{24h}$ post-infection) and the control condition without PZA was used as reference corresponding to 100 % growth. Determination of relative growth index was performed in the absence (pink) or presence (cyan) of v-ATPase inhibitor ConA. Results are displayed as box-plots with individuals' raw data. Black dots were added to highlight the mean of each conditions. **(D)** Determination of PZA $EC_{50}$ in the absence or presence of ConA by performing a four-parameter nonlinear logistic regression of the data displayed in **(A)**. Results are representative from n = 2 biologically independent experiments performed at least in two-three technical replicates. **(E)** Representative confocal fluorescence images of Mtb WT-infected IPSDM for 24 hours and further treated for 72 hours with increasing concentration of PZA. Magnifications display nuclear staining (blue) and Mtb-producing E2-Crimson (red). Scale bar corresponds to 50 µm. Micrographs are representative of 2 independent experiments.
